# Evolution of chromosome arm aberrations in breast cancer through genetic network rewiring

**DOI:** 10.1101/2023.06.10.544434

**Authors:** Elena Kuzmin, Toby M. Baker, Tom Lesluyes, Jean Monlong, Kento T. Abe, Paula P. Coelho, Michael Schwartz, Dongmei Zou, Genevieve Morin, Alain Pacis, Yang Yang, Constanza Martinez, Jarrett Barber, Hellen Kuasne, Rui Li, Mathieu Bourgey, Anne-Marie Fortier, Peter G. Davison, Atilla Omeroglu, Marie-Christine Guiot, Quaid Morris, Claudia L. Kleinman, Sidong Huang, Anne-Claude Gingras, Jiannis Ragoussis, Guillaume Bourque, Peter Van Loo, Morag Park

## Abstract

The basal breast cancer subtype is enriched for triple-negative breast cancer (TNBC) and displays consistent large chromosomal deletions. Here, we characterize the evolution and maintenance of chromosome 4p (chr4p) loss in basal breast cancer. TCGA data analysis showed recurrent deletion of chr4p in basal breast cancer. Phylogenetic analysis of a unique panel of 23 primary tumor/patient-derived xenograft basal breast cancers revealed early evolution of chr4p deletion. Mechanistically we show that Chr4p loss is associated with enhanced proliferation. Gene function studies identified an unknown gene, *C4orf19,* within chr4p, which suppressed proliferation when overexpressed and is a novel member of a PDCD10-GCKIII kinase module, we name as *PGCA1*. Genome-wide pooled overexpression screens using a barcoded library of human open reading frames, identified chromosomal regions, including chr4p, that suppress proliferation when overexpressed in a context-dependent manner implicating network interactions. Together this sheds light on the early emergence of complex aneuploid karyotypes involving chr4p and adaptive landscapes shaping breast cancer genomes.

## Introduction

Breast cancer is a heterogeneous disease comprising several clinical and molecular subtypes. Therapeutic strategies have been devised for patients based on biomarkers, such as hormone (estrogen and progesterone) receptor expression or human epidermal growth factor 2 (HER2) receptor amplification ^1^. However, triple-negative breast cancer (TNBC), constituting 10-20% of all breast cancers, lacks these receptors, thus, lacks precision therapies targeting them and is predominantly treated by chemotherapy. Currently, due to limited therapeutic options TNBC has the most aggressive behavior and worst prognosis (5-year relative survival percent), leading to a large percentage of breast cancer deaths ^2–4^. The basal breast cancer molecular subtype constitutes ∼80% of TNBC and shows a complex mutational spectrum without common oncogenic drivers ^5–7^. Notably basal breast cancers frequently display consistent large chromosomal deletions ^8,9^ that are thought to play an important role in pathogenesis but the consequence of which are poorly understood.

An important hallmark of cancer cells is genomic instability, which generates mutations and chromosome alterations that confer selective advantage on subclones of cells and lead to their growth and dominance in a local tissue ^10,11^. Considerable effort has been invested into identifying which oncogenes, the increased activity, or tumor suppressor genes, the loss of function of which, drive cancer development ^10^. With the advent of genomic technologies, it has become possible to generate a detailed map of genetic changes in cancer ^6,12–14^. Genomic analyses revealed that chromosome arm somatic copy number aberrations are more common than whole chromosome somatic copy number aberrations and certain chromosomal arms are preferentially lost or gained, suggesting that these events are selected because they are advantageous during cancer progression ^15,16^. Evolutionary analyses of Pan-Cancer Analysis of Whole Genomes (PCAWG) data on 38 types of cancer showed that chromosomal arm copy number losses occur early and typically precede gains, indicating their selective advantage in tumor onset and progression ^17^. Recent findings suggest that chromosomal arm aberrations occur in bursts enabling genome diversification and preferential clonal expansion in TNBC ^18^. Although, they have been implicated in some cancers in increasing cell growth ^19^ and evading immune system detection ^20^, the functional consequences of chromosomal arm deletions remain poorly understood.

We previously established that a large deletion on chromosome 5q in basal breast cancer leads to a loss of function of *KIBRA*, encoding a multi-domain scaffold protein, activating oncogenic transcription factors, *YAP/TAZ* ^21^. Chromosome 8p loss in breast cancer alters fatty acid and ceramide metabolism, leading to invasiveness and tumor growth under stress conditions due to increased autophagy, thus contributing to resistance to chemotherapeutic agents ^22^. Another study found that cooperative effects, resulting from genes co-deleted within a region harboring *TP53* on chromosome 17p, lead to more aggressive lymphoma than individual mutations ^23^. Interestingly, a recent study reported differences in specific chromosome arm losses, such as chromosome 3p loss, which positively correlated with immune signatures suggesting that specific chromosomal regions can exert selective pressures rather than overall aneuploidy level^24^.

In this study, we identified chromosome 4p loss as a frequently recurrent chromosome arm loss in the basal subtype of breast cancer and established that this occurred as an early clonal event functionally associated with an enhanced proliferative state. Scanning genes on chr4p by functional assays, we identified genes whose elevated expression suppressed proliferation in human breast epithelial cells. This included an unknown gene, *C4orf19,* within chr4p, we show suppresses proliferation and is a member of the programmed cell death 10 (PDCD10)-germinal centre kinase III (GCKIII) module (which we call *PGCA1*). Genome-wide pooled overexpression screens using a barcoded library of human open reading frames identified chr4p and other chromosomal regions that suppress proliferation when overexpressed in a context-dependent manner. Together this provides new insight into TNBC, a hard to treat cancer, for which the current standard therapeutic options have not significantly changed the overall survival rate.

## Results

### Loss of chromosome 4p is recurrent and functionally significant in basal breast cancer

We investigated the frequency of chromosome arm losses in the breast cancer basal subtype^8^. The three most frequent chromosomal arm losses in basal breast cancer were 8p, 5q and 4p, occurring collectively in ∼65-76% of cases (Fig. 1B& S1A, Table S1). We detected chromosome 4p (chr4p) loss in ∼65% of basal breast cancer cases, which has not been studied previously. Regions of chr4p loss span the entire short arm of chromosome 4 without an apparent minimal deleted region, suggesting that its selective advantage is conferred by the loss of multiple genes residing within the chromosome arm rather than a single tumor suppressor gene (Fig. 1C).

**Figure 1.**
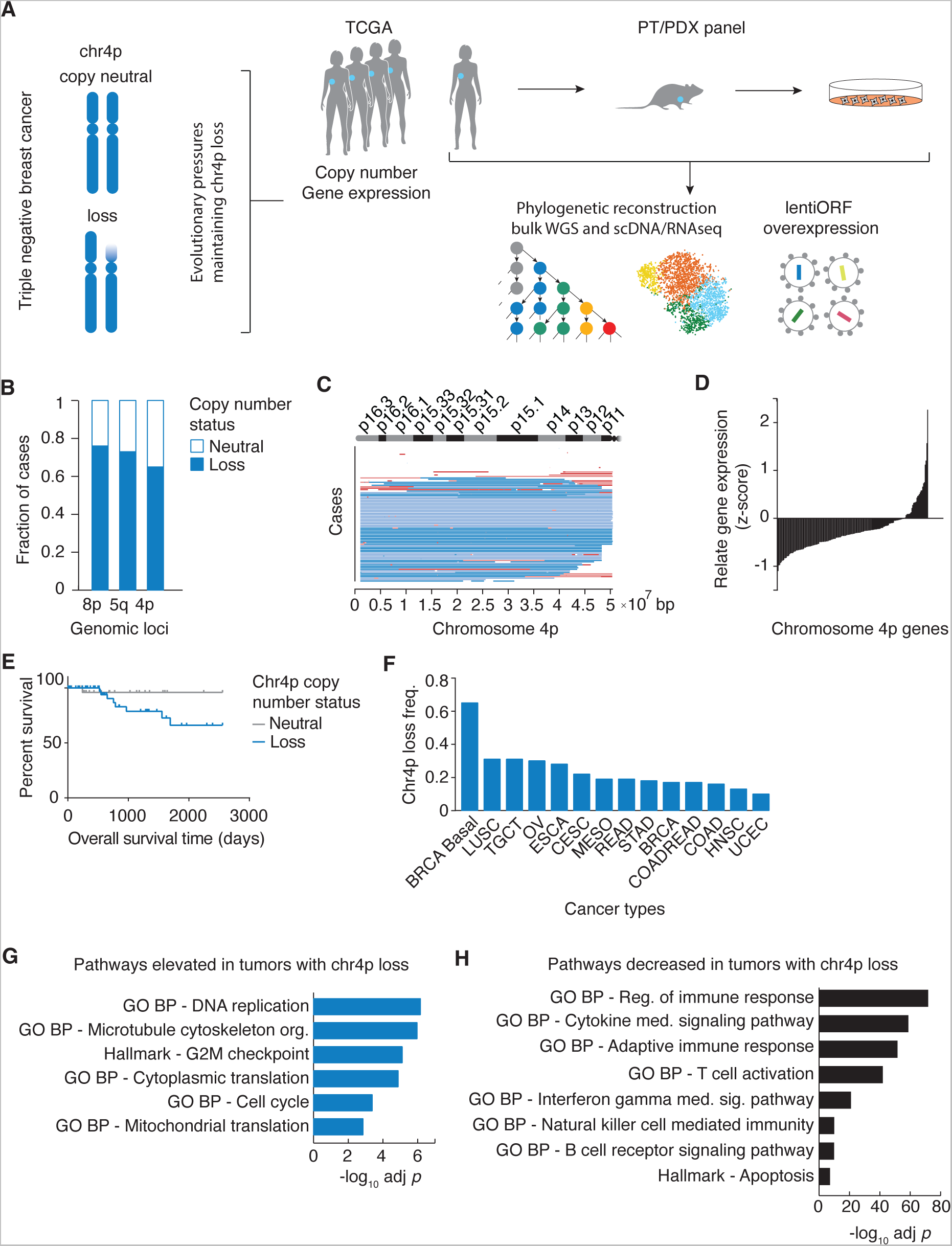
Loss of chromosome 4p in basal breast cancer is recurrent and functionally significant. **(A)** Experimental and analytic pipeline. **(B)** The Cancer Genome Atlas (TCGA) Invasive Breast Carcinoma single nucleotide polymorphism (SNP) array dataset was used to investigate the frequency of chromosome arm losses among the basal subtypes. The three most frequent chromosomal arm losses in basal breast cancer are shown, whereby chr4p loss occurs in ∼65% patients. **(C)** Regions of chr4p loss span a large fraction of the chromosome 4p. Dark blue denotes stringent threshold deletion segmented mean < -0.3, light blue denotes lenient threshold - 0.3 < deletion segmented mean < -0.1, light red denotes lenient threshold 0.1 < deletion segmented mean < 0.3, red denotes stringent threshold deletion segmented mean > 0.3, white denotes copy neutral state. **(D)** TCGA basal breast cancer gene expression dataset was used to show that ∼80% of genes along chr4p decrease in expression upon its copy number loss. **(E)** Overall survival of basal breast cancer patients with copy neutral or deletion status of chr4p, *p* < 0.0997. **(F)** Chr4p copy number status across pan-cancer TCGA datasets. Gene Set Enrichment Analysis (GSEA) showing representative terms that are enriched for genes displaying **(G)** elevated or **(H)** decreased expression due to chr4p loss in TCGA basal breast cancer.

Approximately 80% of genes (133 of 156) within chr4p showed reduced gene expression in patients with chr4p loss, indicating that this loss is functionally significant (Fig. 1D, Table S2). Among the ten genes with the most substantial reduction in gene expression upon chr4p deletion are *SLC34A2* and *RHOH,* that are known tumor suppressor genes in other cancers but have not been previously implicated in breast cancer^25^. Notably the worse survival of cases exhibiting chr4p loss is not due to the enrichment of *TP53* loss of function mutations in chr4p loss since their distribution among groups was not significantly different (Fig. 1E, Methods). Hence, chr4p loss may be clinically prognostic. We also detected statistically significant chr4p deletion in multiple cancer types, including Lung Squamous Cell Carcinoma, Testicular Germ Cell Tumors and Ovarian Serous Carcinoma, indicating that chr4p loss is broadly observed in other cancers (Fig. 1F, Table S3).

Next, we interrogated global transcriptomic changes associated with chr4p loss (Fig. 1G&H, Table S4). Basal breast cancers with chr4p loss showed an elevation of genes with roles in DNA replication (*p* = 6.9×10^-7^) such as *GINS4* encoding a member of the GINS complex, which plays an essential role in initiation of DNA replication and progression of DNA replication forks ^26^ and cell cycle (*p* = 4.1×10^-4^), such as *STK33* encoding a serine/threonine protein kinase, which activates ERK signaling pathway ^27^. Together these terms suggest of a proliferative advantage conferred by chr4p loss likely through a combined effect of expression changes of multiple genes belonging to these pathways. Elevated expression of genes with a role in microtubule cytoskeleton organization (*p* = 1.1×10^-6^) and protein translation (*p* = 1.4×10^-3^), suggest the involvement of cellular plasticity, which enables cancer cell adaptation to stress ^28^. Chr4p loss was associated with decreased expression of genes annotated to positive regulation of immune response, cytokine-mediated signaling, T cell and B cell activation, interferon gamma mediated signaling and natural killer cell-mediated immunity consistent with TNBC displaying immune evasion and poorer outcome. This differential gene expression was not due to general differences in arm-level and chromosome-level copy number changes (*p* = 0.074) (Fig. S2, Table S5) ^20^. These findings support that these global transcriptomic changes are specific to chr4p loss rather than general differences in aneuploidy between tumors and highlight the importance of specific chromosome arm losses in basal breast cancer.

### Chromosome 4p loss is an early event in basal breast cancer evolution

Evolution of basal breast cancer genomes can be reconstructed from somatic mutations detected by whole genome sequencing (WGS)^17^. To understand the evolutionary timing of chr4p loss, we performed phylogenetic reconstruction on bulk WGS of our primary tumor / patient-derived xenograft (PT/PDX) panel of 23 paired samples annotated to PAM50 basal breast cancer subtype for a total of 48 unique samples, which we previously collected^29^. To our knowledge this is the largest available phylogeny for TNBC to date. Aggregated single-sample ordering revealed a typical timing of chromosome arm aberrations and other genetic events (Fig. 2A, Table S6). Coding mutations in *TP53* had a high likelihood of being clonal and thus occurring early in tumor progression, consistent with it being a known driver event in basal breast cancer; as well as preceding whole genome duplication consistent with *TP53* function and recent findings from single-cell, single-molecule DNA-sequencing of 8 human triple-negative breast cancers and four cell lines ^18^. We also observed that chromosomal arm losses occurred before chromosomal gains, and chr17p loss harboring *TP53* occurred with similar timing to *TP53* coding mutations, likely driving biallelic inactivation of *TP53*. Most frequent and clonal chromosomal arm losses included 4p, 17q, 3p, 17p, 15q, 14q, and 5q, the majority of which occurred after *TP53* coding mutations. Even though the median age of diagnosis for this PT/PDX cohort was 54, whole genome duplication occurred ∼6 years prior, around 48 years of age in 16 samples (∼70%) and the most recent common ancestor was observed on average ∼2 years before diagnosis, around 52 years of age (Fig. 2B, Table S6). Unlike early events, there was no apparent subclonal structure since it tended to be distinct among patients, suggesting that even though early events are shared among basal breast cancer patients, late events diverge. These findings are consistent with a recent study on gastric cancer evolution, which showed that *TP53* leads to chromosome arm-level aneuploidy in a temporally preferred order ^30^. Thus, it is important to focus on clonal events for therapeutic application.

**Figure 2.**
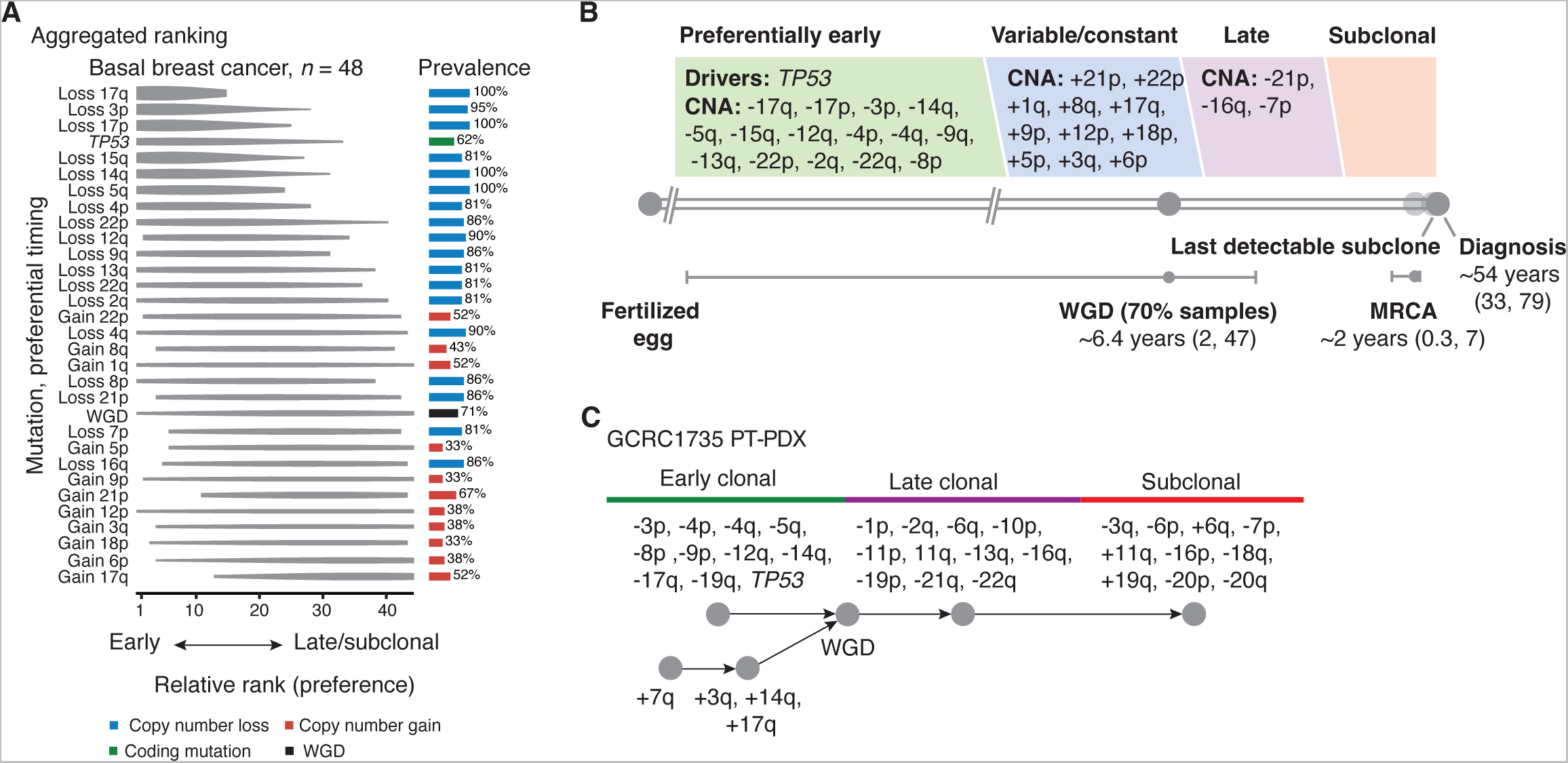
Chromosome 4p loss is an early event in basal breast cancer evolution. Basal breast cancer primary tumor / patient-derived xenograft (PT/PDX) panel was used for the phylogenetic reconstruction. **(A)** Aggregated single-sample ordering reveals typical timing of chromosome arm aberrations. Preferential ordering diagrams show probability distributions revealing uncertainty of timing for specific events in the cohort. The prevalence of the event type in the cohort is displayed as a bar plot on the right. Losses occurring in > 80% and gains occurring in >30% of all cases and evolutionary stages and are depicted. **(B)** Timeline representing the length of time, in years, between the fertilized egg and the median age of diagnosis for breast cancer. Real-time estimates for major events, such as whole genome doubling (WGD) and the emergence of the most recent common ancestor (MRCA), are used to define early, variable, late and subclonal stages of tumor evolution approximately in chronological time. Driver mutations and copy number alterations (CNA) are shown in each stage according to their preferential timing, as defined by relative ordering. **(C)** An example of individual patient (PT/PDX1735) trajectory (partial ordering relationships), the constituent data for the ordering model process.

Since we detected *TP53* coding mutations occurring earlier than chr4p loss and no single known cancer gene was associated with chr4p loss, it is possible that a combination of cancer gene mutations in concert with *TP53* leads to a selective advantage of chr4p loss. To understand if any of the early evolutionary events in our basal breast cancer cohort were already present in preinvasive cancer, we analyzed copy number data of 95 patient samples diagnosed with ductal *in situ* carcinoma (DCIS) in a previously published study^31^. We found that *TP53* coding mutations and clonal losses of 3p, 4p, 5q, 8p, 13q, 14q, 16q, 17p, 17q, 21p, 22q and gains of 1q, 8q and 17q that were present in basal breast cancer were also detected in at least 20% of the DCIS samples indicating that these chromosome arm aberrations are key events in tumorigenesis (Fig. S3A). To understand the selection pressures that maintain chr4p loss, we decided to focus on an individual patient (PT/PDX1735) whose trajectory revealed an early clonal chr4p loss, a clonal *TP53* coding mutation and WGD (Fig. 2C) and for which we established a PDX and PDX-derived cell line. Our scDNAseq analysis of PDX1735 confirmed that chr4p loss was an early event in this basal breast cancer progression (Fig. S3B&C, Table S7).

### Chromosome 4p loss is associated with a proliferative state

To understand the functional effect of chr4p loss, we leveraged single cell RNAseq (scRNAseq) data from PDX1735, established from a basal breast cancer primary tumor, as reported in our previous study (Fig. 3A top left) ^32^. To infer copy number status at a single cell resolution to identify transcriptional programs associated with cells harboring chr4p deletion (Fig. 3), we employed a method, which detects consistent variation in gene expression of consecutive genes across chromosomal regions_33_. To obtain a normal gene expression baseline, we performed scRNAseq on breast tissue samples from two patients undergoing bilateral mammoplasty reduction (Figure S4, Table S8). We computed a z-score relative to the baseline and called copy number aberrations using a Hidden Markov model (HMM) with three states: neutral copy number, loss, and gain. In this manner, we identified four stable ‘communities’, groups of cells with a shared pattern of inferred copy number profiles. Three communities (1-3) harbor chr4p deletion and community 4 harbors a chr4p copy neutral state (Fig. 3A top right, bottom, Table S9). The relatively small size of community 4 (∼ 9%), which is copy neutral for chr4p, is consistent with chr4p loss being an early clonal event, as revealed by our timing analysis (Fig. 2C&S3).

**Figure 3.**
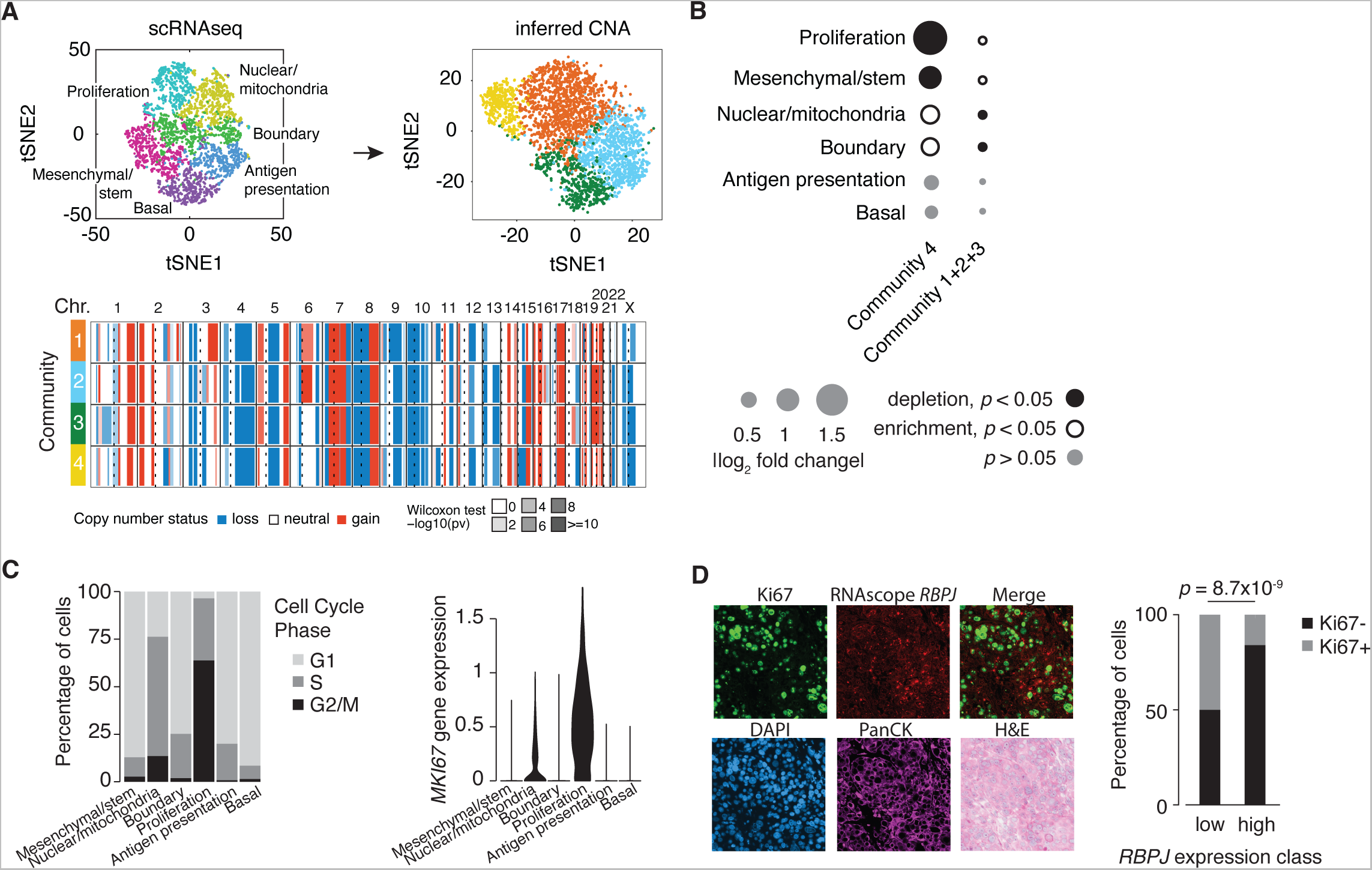
Chromosome 4p loss is associated with a proliferative state. **(A)** Single cell RNA sequencing data of PDX1735 (left panel) from a previous study^32^ were used to infer copy number status (right panel) and displayed using tSNE plots. Groups of cells with shared pattern of gene expression profiles or inferred copy number profiles are colored in the same color. Heatmap (bottom panel) shows the inferred copy number profile of four communities which identified three communities (1-3) harboring chr4p deletion and community 4 harboring chr4p copy neutral state. chr denotes chromosome, dashed line denotes centromere, solid line denotes start/end of chromosome. Loss is in blue, copy neutral is in white, gain is in red. Likelihood of inferred copy number change is represented with Wilcoxon test -log_10_ *p* value, with darker shade reflecting higher confidence. **(B)** Frequency of cells inferred to harbor chromosome 4p deletion or copy neutral state across different cellular clusters with distinct transcriptional programs. The “transcriptional program” category received a count for any combination in which a cell belonged both to a specific inferred copy number community and a specific gene expression cluster. The size of the circle assigned to each “transcriptional program” element reflects the fold increase over the background fraction of all cells in a specific gene expression cluster. Significance was assessed with a hypergeometric test; *p* < 0.05. Solid black circles represent significant depletion; open black circles represent significant enrichment; grey denote no significant change. **(C)** Distribution of cells across (left panel) cell cycle phases (light grey denotes G1 phase, medium grey denotes S phase, black denotes G2/M phase) and (right panel) Ki67 gene expression based on cell clusters with distinct transcriptional programs as shown in A. **(D)** Staining of paraffin embedded fixed tissue section of PDX1735 using a combination of immunofluorescence (IF) for Ki67 (marker of proliferation transcriptional program cell cluster), RNA-*in situ* hybridization (ISH) for *RBPJ* (marker of chr4p), DAPI staining for nuclei, pan Cytokeratin (PanCK) IF for epithelial cancer cells and hematoxylin and eosin (H&E) staining for cancer histology. Significance was assessed by Fisher exact test.

To understand the selection pressures that maintain chr4p loss, we compared the distributions of chr4p copy neutral and deletion communities across different cellular clusters associated with distinct transcriptional programs. We previously described six cellular clusters embedded within PDX1735, which included proliferation, nuclear/mitochondria, antigen presentation, basal, mesenchymal/stem and boundary based on differential gene expression ^32^. As expected, the inferred copy number communities did not overlap with any specific gene expression cluster, since the normalized expression was smoothed using a rolling median approach to reduce the effect of single-gene outliers (Fig. S4B). Thus, cells belonging to chr4p deletion communities (community 1+2+3), which comprised most cells of the PDX1735, did not show a preferential distribution across any cellular cluster (Fig. 3B). However, cells belonging to the chr4p copy neutral community (community 4) were significantly strongly depleted in the proliferation and to a lesser extent in the mesenchymal cluster (Fig. 3B). The proliferation cluster was characterized by proliferative cells since a large proportion of cells in this cluster (∼95%) were cycling and exhibited inferred G2/M and S cell cycle states based on the relative gene expression of G1/S and G2/M gene sets ^34^ as well a high expression of cell cycle genes, such as *MKI67* (Fig. 3C, Table S9).

To functionally validate the findings from scRNAseq data, we performed immunofluorescence staining combined with RNA-*in situ* hybridization (RNA ISH). The staining of a paraffin-embedded fixed tissue section of PDX1735 used a combination of immunofluorescence for Ki67 and RNA ISH for *RBPJ*. *RBPJ* was selected as a marker of chr4p copy number state because of the availability of a probe for RNA ISH and a consistent gene expression difference in our basal breast cancer PT/PDX panel between chr4p copy neutral and deletion samples. This analysis revealed that there was an inverse relationship between Ki67 abundance and *RBPJ* gene expression. Breast cancer cells with chr4p deletion and thus low *RPBJ* expression showed a high abundance of Ki67 and thus were more proliferative than chr4p copy neutral cells (Fisher exact test, *p* = 8.7×10^-9^) (Fig. 3D). Together these findings suggest that chr4p loss confers on basal breast cancer cells a proliferative advantage.

### Suppression of proliferation by overexpression of chromosome 4p genes is context-dependent

To determine if chr4p deletion in basal breast cancer is selected due to a proliferative advantage, we tested whether the overexpression of genes within this region elicits a proliferation defect. Chr4p copy neutral normal breast epithelial cell line, MCF10A, chr4p copy neutral basal breast cancer PDX-derived cell line, GCRC1915, or chr4p deletion basal breast cancer PDX-derived cell line, GCRC1735 were used to generate stable cell populations overexpressing candidate chr4p genes using lentivirus-mediated integration of constructs from the human ORF collection (Fig. 4A) ^35^. The candidate genes resided within a high confidence chr4p deletion region in GCRC1735 according to whole exome sequencing (WES) from our previous study ^32^, encompassing about half of the chromosome arm and which contained 30 genes, for about half of which our collection contained lentiORF overexpression vectors.

**Figure 4.**
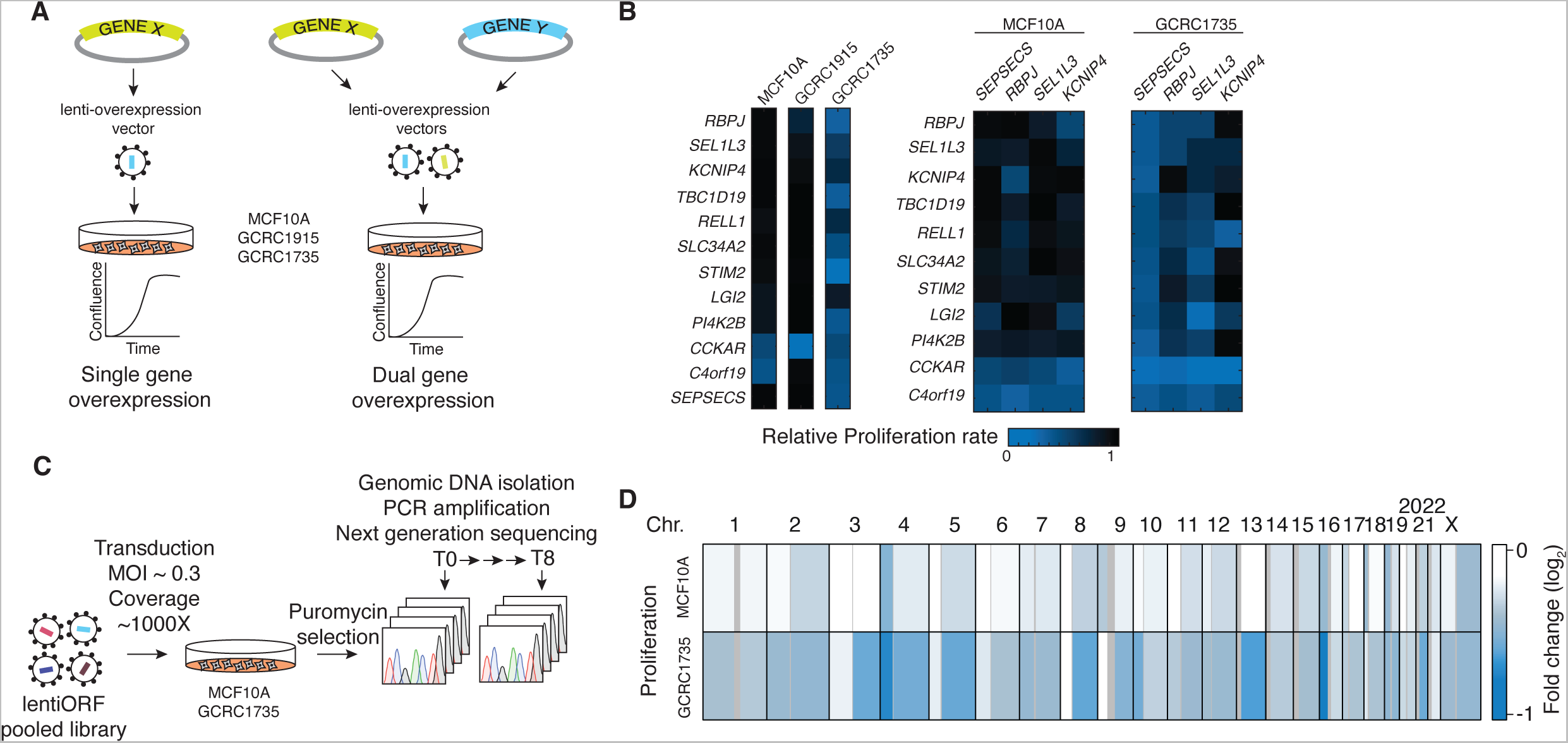
Overexpression of chromosome 4p genes leads to context-dependent suppression of proliferation. **(A)** A schematic of gene overexpression strategy. Chr4p copy neutral normal breast epithelial cell line (MCF10A), chr4p copy neutral basal breast cancer patient derived xenograft (PDX) -derived cell line (GCRC1915) or chr4p deletion basal breast cancer PDX-derived cell line (GCRC1735) were used to generate stable cell populations using lentivirus-mediated integration. The resulting mutant cell line populations overexpressed candidate genes residing within chr4p region, which is found in a high confidence chr4p deletion region in GCRC1735 according to whole exome sequencing (WES) from a previous study ^32^. Lentiviral constructs for single and dual gene overexpression were obtained from the ORF collection^35^. **(B)** Chr4p gene overexpression confers a proliferation defect in GCRC1735 with chr4p loss but not in chr4p copy neutral cell lines (MCF10A, GCRC1915). Dual overexpression exacerbates the proliferation defect observed in GCRC1735. Blue denotes proliferation defect, black no change relative to control. **(C)** Schematic of the genome-wide pooled lentiORF overexpression screen. MCF10A and GCRC1735 cell lines were transduced with pooled lentiORF library at multiplicity of infection (MOI) of 0.3. After puromycin selection cells were maintained for 6-8 doublings and 1000x coverage was maintained at each step of the experiment. Next generation sequencing (NSG) was used to capture barcode abundance which served as a proxy for cell growth rate. **(D)** Genome-wide pooled lentiORF overexpression screen revealed genomic regions with context dependent suppression of proliferation. For example, there was a growth defect in GCRC1735 and not in MCF10A, including chr4p and 13q that are deleted in GCRC1735 and not in MCF10A. Blue denotes proliferation defect, black no change relative to T0 control.

Surprisingly, overexpressing a large fraction of chr4p genes suppressed proliferation in a context-dependent manner, whereby proliferation suppression was only observed in a basal breast cancer PDX-derived cell line which is deleted for chr4p, PDX1735, and not in cell lines that were chr4p copy neutral, MCF10A or GCRC1915. Of note GCRC1915 displays LOH within chr4p it is copy neutral with no change in chr4 copy number (Fig. 4B left). This extent of suppression was further exacerbated when two random genes within chr4p were overexpressed (Fig. 4B right). The context-dependency of suppression of proliferation of chr4p genes is likely not due to *TP53* mutation status since both GCRC1915 and GCRC1735 harbor a coding mutation in *TP53*. On the other hand, GCRC1735, unlike GCRC1915, harbors a germline *BRCA1* mutation^29^, clonal losses chr1p, chr19p, chr19q, chr21q, chr22q and clonal gains chr3q, chr7q, chr14q, chr17q (Table S6), which may underlie the context dependent suppression of proliferation of chr4p gene overexpression observed in GCCR1735 and not in GCRC1915. Since both chr4p and chr22q losses are often early clonal events in basal breast cancer (Fig. 2B), it is possible that their interaction rewires chr4p maintaining it in a deletion state by selecting against chr4p amplification. Hence, the observed context-dependent suppression of proliferation may be due to a genetic interaction with another genetic aberration, which rewires the genetic network sensitizing chromosome 4p region to overexpression and thus maintaining it in a deletion state. Further studies in isogenic model systems should be conducted to test which combinations of genetic events interact with chr4p loss to confer a proliferative advantage, since it was previously found that individual chromosome arm losses lead to growth defects ^24^.

To determine whether the context-dependent suppression of proliferation was specific to chromosome 4p and to identify other such regions, we conducted a genome-wide overexpression screen using pooled TRC3 LentiORF collection as previously described (Fig. 4C, Table S11) ^36^. Two cell lines with different copy number states of chr4p, MCF10A (chr4p copy neutral) and GCRC1735 (chr4p deletion), were transduced with pooled lentiORF library at multiplicity of infection (MOI) of 0.3 to ensure one integration event per cell. After puromycin selection cells were maintained for 6-8 doublings and 1000x coverage was maintained at each step of the experiment. Next generation sequencing (NSG) was used to capture barcode abundance, which served as a proxy for cell growth rate. Genome-wide pooled lentiORF overexpression screen uncovered genes that suppressed proliferation in both cell lines and were previously identified as STOP genes in another study, such as epithelial tumor suppressor *ELF3*, transcription factor *EBF1* and a DNA repair protein *RAD51* ^19^. The screen also revealed regions that suppressed proliferation in a context-dependent manner. The context-dependent regions that suppressed proliferation when overexpressed in GCRC1735 but not in MCF10A included chr4p and 13q. These regions were also deleted in GCRC1735 and not in MCF10A (Fig. 4D), suggesting that this mode of selection is not specific to chr4p loss and likely exerts the selection pressure early in tumor progression since our evolutionary timing analysis revealed that both are clonal events (Fig. 2C). These observations suggest that the dosage of these genes exerts a selection pressure to maintain this chromosomal region deleted in a specific genomic context of basal breast cancer.

### Overexpression of C4orf19 suppresses proliferation and reveals an interaction with PDCD10-GCKIII kinase module

We observed that *C4orf19* suppressed proliferation when overexpressed in multiple contexts, such as MCF10A and GCRC1735 cells (Fig. 4B). To understand the biological role of *C4orf19*, we analyzed the sequence of the protein it encodes. C4orf19 is an uncharacterized protein 314 amino acids in length, which has orthologs in mouse and rat according to the Alliance of Genome Resources ortholog inference ^37^. Functional analysis of its protein sequence using InterPro revealed that it belongs to the protein family domain unknown function DUF4699 and is predicted to contain two consensus disorder regions (36-142, 267-291). We mined BioGRID ^38^ for previously identified protein-protein interactions: high throughput methods, such as affinity capture-MS and yeast two-hybrid both revealed programmed cell death 10, PDCD10 (Fig. 5A) ^39,40^. Additionally, all three members of the germinal center kinases (GCK-III) subfamily, STK24, STK25, STK26 that directly interact with PDCD10 in a mutually-exclusive heterodimer ^41^ were reported in the affinity capture-MS method_40_. We heterologously expressed C4orf19-v5 and 3xFLAG-PDCD10, 3xFLAG-STK25 and 3xFLAG-STK26 in MCF10A cells (Fig. 5B). Co-immunoprecipitation assay using Anti-V5 for pull-down showed the presence of 3xFLAG-PDCD10, 3xFLAG-STK25 and 3xFLAG-STK26, confirming previously identified high-throughput interactions, indicating that C4orf19 interacts with PDCD10 and its associated GCK-III kinases (Fig. 5C).

**Figure 5.**
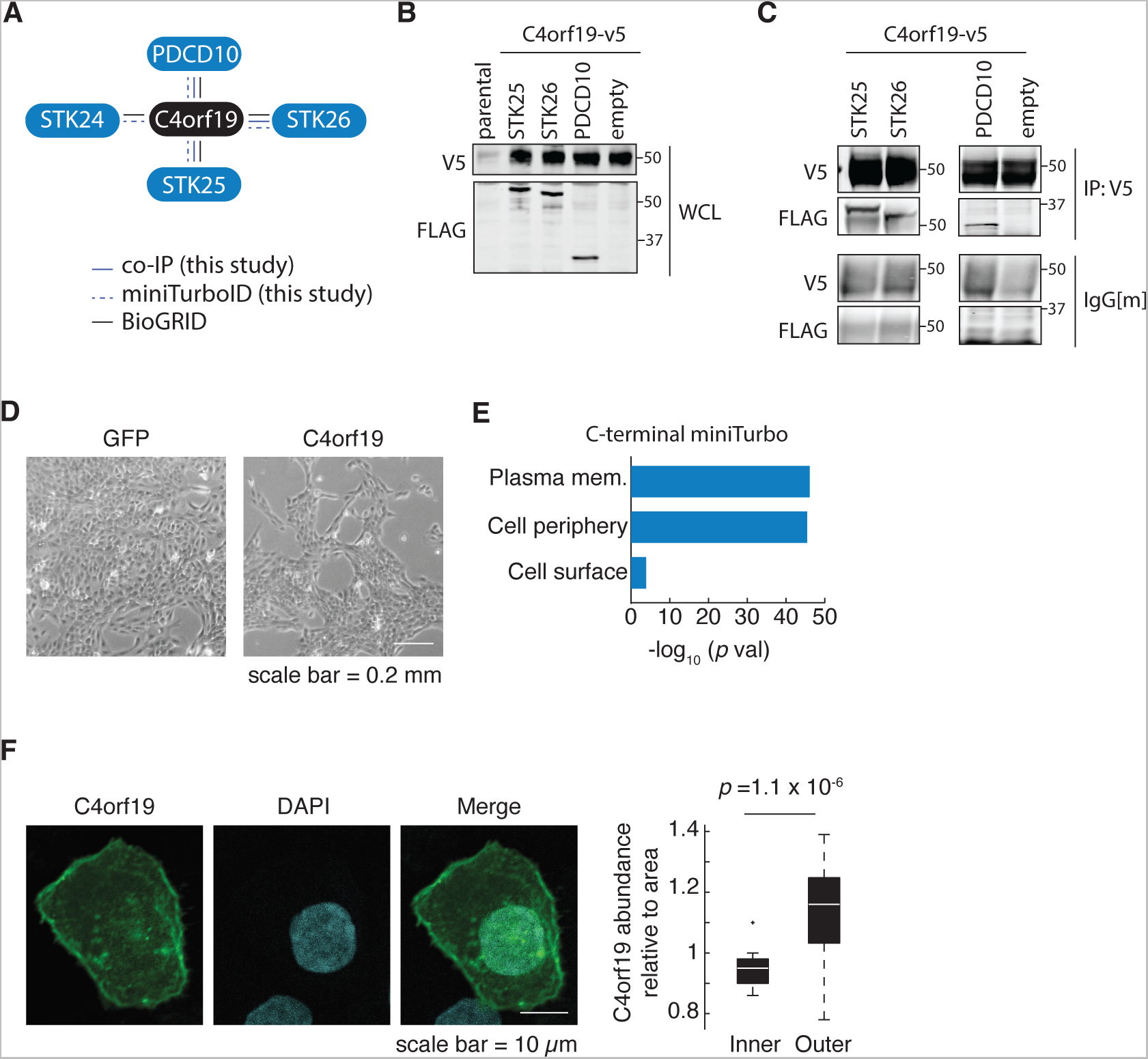
**C4ORF19** (PGCA1) **is associated with the PDCD10-GCK-III module**. **(A)** Summary of co-IP assay results from (B), miniTurbo ID conducted in MCF10A cells expressing C4rf19-miniTurbo and literature curation using BioGRID. **(B)** Western blot using whole cell lysate shows heterologous expression of C4orf19-v5 and PDCD10-GCK-III module 3xFLAG-PDCD10, 3xFLAG-STK25 and 3xFLAG-STK26 in MCF10A cells. **(C)** Co-immunoprecipitation assay using Anti-V5 for pull-down shows the presence of 3xFLAG-PDCD10, 3xFLAG-STK25 and 3xFLAG-STK26 in MCF10A cells indicating that C4orf19 interacts with PDCD10 and associated GCK-III kinases. **(D)** Brightfield microscopy images of MCF10A cells overexpressing GFP control or C4orf19 48 hrs post induction with doxycycline. **(E)** Analysis of gene ontology molecular function (GO MF) of proteins in proximity to C4orf19 from miniTurbo ID from (C) shows enrichment of proteins at the plasma membrane suggesting that C4rof19 is localized to the cell periphery. **(F)** Immunofluorescence of C4orf19 indicates that its subcellular localization is at the cell periphery. Signal intensity relative to area of the inner or outer cell section indicates that C4orf19 abundance is higher at the cell periphery, n = 27. Significance was assessed using Wilcoxon rank sum test.

To gain more insight into the biological role of C4orf19, we performed proximity-dependent biotinylation of proteins coupled to mass spectrometry (miniTurboID) ^42^ to reveal the comprehensive physical neighbourhood in which C4orf19 resides. MiniTurbo biotin ligase was fused to C4orf19 and expressed in MCF10A (alongside negative controls; bait expression was verified by western blotting), and biotinylation of proximal proteins was induced by the addition of biotin (Fig. S5A&B). Biotinylated proteins were recovered by streptavidin-affinity chromatography and identified by mass spectrometry. Reduction in proliferation was observed 48 hr after induction of *C4orf19* overexpression with doxycycline (Fig. 5D). Using the SAINTexpress computational tool, we identified 370 high-confidence (Bayesian FDR < 5%) proximal interactors, which included PDCD10, STK24, STK25 and STK26 (Fig. 5A, Table S10). The analysis of gene ontology molecular function (GO MF) of C4orf19 proximal interactors showed enrichment of proteins at the plasma membrane, suggesting that C4orf19 is localized to the cell periphery (Fig. 5E). The subcellular localization of C4orf19 at the cell periphery was further validated by immunofluorescence of C4orf19. The signal intensity of C4orf19 relative to area of the inner or outer cell section indicates that C4orf19 abundance is higher at the cell periphery although it was not uniformly distributed (Fig. 5F). This finding is consistent with Human Protein Atlas, which showed that across tissues C4orf19 is at the cell periphery in some cancers, such as breast lobular carcinoma and RT4 cells derived from a urinary bladder transitional cell papilloma ^43,44^. Similarly, the Alliance of Genome Resources computationally predicted its localization to cell junctions based on GO annotation and orthology_37_. We did not observe C4orf19 in the nucleus, which is provided as a secondary predicted localization of this protein, which may be due to cell type specificity.

These findings support that C4orf19 is associated with a subset of PDCD10 and GCK-III kinases localized to the cell periphery, and we propose to rename *C4orf19* to *PGCA1* (*P*DCD10-*GC*K-III Kinase *A*ssociated). *STK24* resides on chr13q and *STK25* on chr2q, both of which show a deletion state in GCRC1735 and suppress proliferation when overexpressed (Fig. 4D), suggesting of copy number change as a selection for stoichiometric balance of the members of the PDCD10-GCKIII kinase module. Overall, the deletions of chr13q and chr4p are both early clonal events in the basal PT/PDX panel highlighting a potential importance of the stoichiometric balance of the PDCD10-GCKIII kinase module as an additional common evolutionary mechanism in this breast cancer subtype.

## Discussion

This study identified chromosome 4p loss as a frequently recurrent chromosome arm loss in the basal breast cancer molecular subtype, affecting ∼65% of TNBC patients. Our data indicate that chr4p loss is functionally significant. It is associated with reduced expression of most genes within the region and poor prognosis. An evolutionary timing analysis revealed that chr4p loss is an early clonal event and was already detected in DCIS, which is the most common form of preinvasive breast cancer. Through multiple approaches we showed that the deletion of chr4p is associated with enhanced proliferation. Targeted single and dual gene overexpression assays of genes within chr4p uncovered *C4orf19,* which suppressed proliferation and was identified to be associated with the PDCD10-GCKIII kinase module, and we propose to rename this gene to *PGCA1*. However, most genes within chr4p suppressed proliferation when overexpressed in a context-dependent manner associated with chr4p deletion. Genome-wide pooled overexpression screens identified other chromosome arms whose suppression of proliferation was context-dependent and associated with copy number loss. Our findings support chr4p loss confers a proliferative advantage in basal breast cancer, and multiple genes within chr4p suppress proliferation when overexpressed in chr4p loss but not copy-neutral cells. Together this provides a unique understanding of the early emergence of complex aneuploid karyotypes involving chr4p and the adaptive landscapes shaping breast cancer genomes.

We found that chromosome arm losses are hemizygous, likely due to the presence of core essential genes residing within them (Fig. S1A) ^45,46^. The apparent lack of a clear minimal deletion region of chr4p in basal breast cancer suggests that this event is not driven by the loss of a single tumor suppressor gene but rather by multiple genes and/or genetic elements at multiple, spatially separated loci whose deletion together yields a proliferative advantage. While there were four tumor suppressors genes within chr4p, none of them has been implicated in breast cancer^25^. These include *SLC34A2,* which encodes a pH-sensitive sodium-dependent phosphate transporter ^47^ and *N4BP2*, which encodes 5’-polynucleotide kinase, playing a role in DNA repair, which have been implicated in lung cancer ^48^. *RHOH* encodes a member of the Ras superfamily of guanosine triphosphate (GTP)-metabolizing enzymes and has been implicated in non-Hodgkin’s lymphoma ^49^. *PHOX2B* is a transcription factor involved in neuroblastoma ^50^. In contrast to *PHOX2B*, which is recessive, the tumor suppressive effects of *SLC34A2* and *RHOH* are dominant ^25^, indicating that the perturbation of one copy of these genes is sufficient to contribute to carcinogenesis and their combined effect due to chr4p loss may be providing a selective advantage in TNBC. Surprisingly, the expression of several genes is elevated, likely due to the loss of *cis*-transcriptional repressors. These include *UCLH1*, which is a ubiquitin hydrolase previously shown to be highly expressed in metastatic estrogen receptor negative (ER–) and triple negative breast cancer subtypes ^51^.

Simultaneous gene expression silencing using RNAi of multiple combinations of up to three genes along chromosome 8p inhibited tumorigenesis in a mouse model of hepatocellular carcinoma (HCC) indicating that multiple TSGs show a greater capacity to promote tumorigenesis than individual genes ^52^. The effect of chromosome 3p loss in lung cancer is also attributed to an alteration in a combination of genes ^24,53^. The apparent lack of a minimal region in a large chromosomal aberration has been also previously observed in human embryonic stem cells and induced pluripotent stems cells when screening for genetic changes occurring in cell culture to evaluate their tumorigenicity, which reported a recurrent gain of chromosome 1, 12 and 17 without any frequently repetitive minimal amplicon ^54^. The gain of chromosome 12 or its short arm 12p has been also shown to arise rapidly during the reprogramming process and is associated with gene expression changes indicating its functional significance in conferring selective growth advantage ^55^.

The functional significance of chr4p loss as assessed by the reduction in gene expression of a large portion of genes within the region, as well as the association of the chromosome arm loss with a proliferative state and mesenchymal state, are consistent with previous observations related to chr3p loss, whereby hallmark sets of cell cycle and epithelial-mesenchymal transition genes were upregulated when chr3p was lost ^24^. Chr4p loss in basal breast cancer was also associated with a decrease in immune signature suggesting that specific partial aneuploidy of chr4p rather than general differences in aneuploidy between tumors may be important for immune system evasion, which were previously reported in a pan-cancer analysis ^20^. This is especially important since it was previously shown that triple-negative breast cancer with an “immune-cold” microenvironment characterized by the absence of CD8+ T cells in the tumor resulted in poor outcomes ^56^. The finding of chromosome arm specific aneuploidy exerting distinct effects on the immune system is likely due to different immure-related genes residing there which has recently emerged, such as a finding that chr3p loss is associated with increased immune activity ^24^.

It is thought that changes in the copy number of specific genes due to large chromosomal variants lead to increased cell fitness. We previously established in a murine model that 5q loss of heterozygosity leads to a loss of function of *KIBRA*, which encodes a multi-domain scaffold protein that inhibits oncogenic transcriptional co-activators YAP/TAZ that mediate mechanotransduction signals ^21^. Chr5q also harbors *RAD17*, *RAD50* and *RAP80* genes that are important for *BRCA1*-dependent DNA repair, and their loss impairs *BRCA1*-pathway function critical for DNA damage control, contributing to increased genomic instability and cancerous phenotype ^57^. Since noncoding genes including lncRNA and miRNA reside within chr4p, it is possible that their loss leads to overexpression of certain target genes contributing to a proliferation state. The observation that chr4p loss is prevalent in *HER2* amplified group but is not associated with decreased survival (Fig. S1A&B), likely indicates that *HER2* amplification, a key event in these tumors, masks the effect, whereby the double mutant carrying both chr4p loss and *HER2* amplification resembles the more extreme phenotype of the single mutant *HER2* rather than the combined effect of both aberrations. The timing analysis showed a preferred order for chromosome arm-level CNA in basal breast cancer as reported in a study on gastric cancer organoids ^30^. In addition, in both studies, these evolutionary early alterations were observed in similar chromosome arms, such as the loss of chr3p, chr4p and chr4q and suggest convergent mechanisms for regulating tumor emergence in gastric and breast cancer, which may be applicable to other cancers. Finally, the finding that chr4p loss along with other early evolutionary events in breast cancer are already detected in DCIS, which is the most common form of preinvasive breast cancer, indicates the key role that these chromosomal aberrations play in tumor initiation and progression.

Our study revealed that *PGCA1* (*C4orf19*) is physically associated with PDCD10 and a subfamily of GCK-III: STK24, STK25, and STK26. It is possible that PGCA1 is a direct binder of PDCD10 since the likelihood of an interaction between two human proteins in a yeast two-hybrid assay when at least one is not nuclear, PDCD10, is very high to be direct ^39^. PGCA1 is likely excluded from the STRIPAK complex, a large multiprotein assembly, the striatin-interacting phosphatase and kinase (STRIPAK) complex, which was initially characterized in HEK293 cells ^58^, since there is no evidence of striatins in the PGCA1 miniTurboID data or other publicly available protein-protein interaction data. This finding further supports the direct binding of PGCA1 to PDCD10 and suggests that PGCA1 competes with striatins for binding to PDCD10. The interface that is mediating this interaction between PGCA1 and PDCD10 may thus be the same as the interface required for binding of striatins to PDCD10 or other partners which was previously shown to be occurring in a mutually exclusive manner ^41,59^. Thus, we propose that PGCA1 is a protein that can tether the GCK-III kinases through the PDCD10 adapter to the plasma membrane bridging the GCK-III kinases to a substrate in this locale.

*PDCD10* is also known as *CCM3* is a causative gene of cerebral cavernous malformation, a neurovascular disease that is characterized by vascular malformations ^60^. In addition to interacting with and controlling signals emanating from the CCM2/CCM1 pathway in cytoskeletal organization, PDCD10 also regulates STRIPAK, and potentially other pathways also implicated in vascular integrity ^60^. PDCD10 is also pro-apoptotic and controls the cell cycle, entry into senescence, apoptotic response to oxidative damage, inflammation and DNA damage repair as well as cell migration ^60^. Its many functions are thought to be enabled by its multiple subcellular localizations, including cell-to-cell junctions and the Golgi apparatus. Our findings suggest that since PGCA1 is at the cell periphery, it likely interacts with PDCD10 at the cell-to-cell junctions. PDCD10 heterodimerizes with GCK-III kinase subfamily and modulates cell migration by regulating Golgi assembly which is mediated by its interaction with STK25 ^61^, regulates exocytosis through its interaction with STK24, which when lost results in oxidative damage and dismantling of the adherens junctions ^62^, as well as maintains ion homeostasis through its interaction with STK26 ^63^. Thus, PGCA1, through its interaction with PDCD10, may tether GCK-III: STK24, STK25 and STK26 to the membrane. This could affect diverse processes that decrease cell proliferation when *PGCA1* is overexpressed and contribute to the proliferation and mesenchymal transcriptomic signatures observed in cell populations with chr4p loss. The similarity of the context-dependent suppression of proliferation of chr4p genes to chr13q, a region deleted in ∼45% basal breast cancers, which also harbors a PDCD10 heterodimerization partner, STK24, highlights the important role of PDCD10-kinase module stoichiometric balance in exerting selection pressures on copy number evolution of breast cancer genome.

It has been previously proposed that the cancer genome is shaped by sensitivity to a change in gene dosage caused by chromosome arm loss or gain ^64^. The gene dosage balance hypothesis postulates that the balance in the ratio of oncogenes to tumor suppressor genes exerts a selective pressure on the cancer cell. Thus, chromosome arm loss will be favoured if the number of tumor suppressor genes is higher than oncogenes and *vice versa* in the case of chromosome arm gain. Our observation of context dependent suppression of proliferation of a frequently lost chromosome arm suggests that additional mechanisms exist that maintain cancer cells with specific chromosome arm losses. A recent pan-cancer evolutionary study noted recurrent early genetic events and the broadening of this set in later stages, suggesting a preference for these genomic changes in early tumor evolutionary stages and potential genetic interactions that constrain the evolution ^17^. This is consistent with previous studies that suggested that despite aneuploidy resulting in a growth disadvantage due to proteotoxic and metabolic stress, it may lead to increased selective pressure on cells to acquire growth-promoting genetic alterations.

Aneuploid genomes are inherently plastic and were shown to evolve an improved fitness state by acquiring additional chromosomal gains and losses ^24,65^. Since co-occurring aneuploidies are often observed in stem cell cultures, tumors and yeast cells, genetic interactions between aneuploidies have been recently suggested to be involved in cancer genome evolution ^66^.

Specifically, in yeast lethality associated with an additional chr VI (which harbors *TUB2*) can be suppressed by an extra copy of chr XIII (carries *TUB1* and *TUB3*), thus supplying a cell with extra alpha tubulin rescuing the stoichiometry imbalance caused by increased level of *TUB2* ^67^. Another study in yeast showed that an additional copy of chr XIII exhibited a growth advantage in combination with chr I gain but a severe growth defect with chr X gain indicating that positive and negative genetic interactions among aneuploidies play a key role in the emergence of complex aneuploid karyotypes ^68^.

The suppression of proliferation of chr4p genes when overexpressed suggests a mechanism of negative selection of chr4p gain in cancers. We showed that chr4p loss was broadly observed across multiple cancer types. The recurrence of chr4p loss in ovarian serous carcinoma is not surprising since this cancer type shares many molecular features and was suggested to have similar etiology to basal breast cancer ^8^. A recent pan-cancer analysis of cancer aneuploidy similarly detected chr4p loss in squamous, gynecological and gastrointestinal tumors ^24^. Since chr4p loss contains multiple genes that promote tumorigenesis when co-deleted, their simultaneous loss may result in vulnerabilities that cannot be identified by studying single genes and thus could provide potential novel therapeutic avenues for patients with triple-negative breast cancer and other cancers.

## Acknowledgements

We thank Anie Monast and Virginie Pilon for assistance with animal experiments, Valentina Munoz Ramos for lab organization and management, Margarita Souleimanova and Nicholas Bertos with handling mammoplasty reduction surgery tissue, Chris Law and the Center for Microscopy and Cellular Imaging at Concordia University for guidance and help with the image analysis. Single cell RNA and DNA sequencing library preparation and sequencing was performed by Yu Chang Wang at the McGill Genome Centre, confocal microscopy was performed at the Centre for Microscopy and Cell Imaging, Concordia University and Advanced BioImaging Facility, McGill University. Histology sections were prepared and imaged in the Histology Innovation Platform, lentiviral ORF collections were obtained from McGill Platform for Cell Perturbation and flow cytometry was conducted at the Flow Cytometry Core Facility, Goodman Cancer Institute. Proteomics was performed at the Network Biology Collaborative Centre at the Lunenfeld-Tanenbaum Research Institute (supported by the Canada Foundation for Innovation, the Ontario Government, Genome Canada, and Ontario Genomics [OGI-139]). The breast tissue bank at McGill University (from which PDXs and cell lines were generated) is supported by the Réseau de Recherche en Cancer of the Fonds de Recherche du Québec-Santé and the Québec Breast Cancer Foundation (to M.P.). This work was supported by the Cancer Research Society (M.P.), the Canadian Institutes for Health Research (FDN 143301 to A.-C.G. and FDN 143281 to M.P.), the Terry Fox Research Institute (to A.-C.G.), Genome Canada Genomic Platform grant (CFI 33406 and 40104 to J.R.; CFI MSI 35444 to J.R.) and Worldwide Cancer Research (16-0402 to M.P.). T.M.B., T.L. and P.V.L. were supported by the Francis Crick Institute which receives its core funding from Cancer Research UK (CC2008), the UK Medical Research Council (CC2008), and the Wellcome Trust (CC2008). For the purpose of Open Access, the authors have applied a CC BY public copyright license to any Author Accepted Manuscript version arising from this submission. P.V.L. is a Winton Group Leader in recognition of the Winton Charitable Foundation’s support towards the establishment of The Francis Crick Institute. P.V.L. is a CPRIT Scholar in Cancer Research and acknowledges CPRIT grant support (RR210006). Additional support was provided by Fonds de Recherche du Quebec-Sante Postdoctoral Fellowship to E.K., L’Oréal Canada Women in Science Research Excellence Fellowship to E.K., Canadian Institutes of Health Research Banting Postdoctoral Fellowship to E.K. and Cancer Research Society Scholarship for the Next Generation of Scientists Postdoctoral Fellowship to E.K., and CIHR Doctoral Fellowship to K.T.A. T.M.B. is supported by a PhD fellowship from Boehringer Ingelheim Fonds.

## Author contributions

Conceptualization: E.K., and M.P.; Methodology and investigation: E.K., K.T.A., P.P.C., M.S., D.Z., G.M., H.K., A.M.F. and J.R.; Formal analysis: E.K., T.M. B., T.L., J.M., K.A., G.M., A.P., Y.Y., J.B., R.L., M.B., M.C.G., A.O., Q.M., C.K., S.H., A.C.G., J.R., G.B., P.V.L., and M.P.; Resources: C.R.M., P.G.D.; Writing – original draft: E.K., and M.P.; Writing – review and editing: E.K., T.B., T.L., J.M., K.A., P.P.C., C.M., J.B., H.K., R.L., M.B., A.M.F., A.O., Q.M., C.L.K., S.H., A.C.G, J.R., G.B., P.V.L., and M.P.; Supervision: M.P.; Funding acquisition: A.C.G., J.R., P.V.L. and M.P.

## Declaration of interest

The authors declare no competing interest.

## Methods

### Isolation of Normal Breast Epithelial Single Cell Suspension

All tissue was collected with informed consent under REB-approved protocols at the McGill University Health Centre. Two patients age of 46 and 18 years old undergoing bilateral mammoplasty reduction due to hypertrophy of the breast with diagnosis and/or management at McGill University Health Centre, Montreal, QC, Canada were recruited for this study. 2,000– 3,000 mm^3^ surgically removed breast epithelial tissue was harvested and kept on ice in transport medium: RPMI 1640, 50 μg/mL gentamycin, 100 U/mL Pen/Strep, 2.5 μg/mL Fungizone until sample processing. The tissue dissociation was achieved as previously described ^33^. Briefly, the tissue fragment was minced in ∼1 mL cold DMEM and MidiMACS Starting Kit (LS) was used as per manufacturing instructions. Minced tissue was collected in a sterile gentleMACS C tube and enzyme A, enzyme R and enzyme H were added from the Tumor Dissociation Kit. GentleMACS Octo Dissociator with Heaters, program was run (1 h, mild speed, 37 °C) to begin the mechanical and enzymatic digestion process. The mix was incubated on ice for 3 min to allow for gravity sedimentation and the oily layer was aspirated. 3 ml of cell suspension was run through 70 μm strainer and collected. The remaining undigested tissue fragments were loaded into the gentleMACS Octo Dissociator, program (1 min, high speed, room temperature) to continue the mechanical and enzymatic digestion process. The cell suspension was passed through 70 μm strainer and collected. The strainer was washed with 10 mL PBS and combined with the filtrate, centrifuge for 10 min, 1,500 rpm. The cell pellet was resuspended in 500 μL Complete DMEM and Trypan Blue staining was used to quantify cell number and viability. Red blood cells were removed by aspirating the supernatant and adding 3 mL ACK lysing buffer and incubating at room temperature for 5 min. Then, 7 mL PBS were added and the suspension was centrifuged at 1,200 rpm for 4 min. The cell pellet was then resuspended in 500 μL complete DMEM. Single cell RNA sequencing was conducted if the cell viability exceeded 60%.

### Cell culture

MCF10A cell line was obtained from the ATCC and cultured in DMEM/Ham’s F12 medium, 20 ng/ml hEGF, 100 ng/ml cholera toxin, 10 μg/ml bovine insulin, 500 ng/ml hydrocortisone, 5% horse serum (HS), 50 μg/ml gentamicin. HEK293T cell line was cultured in DMEM with 10% fetal bovine serum (FBS).

Patient-derived xenograft derived cell lines (GCRC1735, GCRC1915) were isolated from the respective PDXs. PDX tumor fragments were minced and digested as previously described ^33^. Cancer epithelial cells were established using a Conditional reprogramming protocol as previously described ^69^. Briefly, after tumor fragments digestion, single-human epithelial cancer cells were transferred to a dish containing lethally irradiated 3T3-J2 cells (1 × 106 cells) and cultured with F-media (DMEM (Gibco) and F-12 Nutrient Mixture (Ham) (Gibco-) (1:4), 5% FBS (Life Technologies), 0.4 ug/mL Hydrocortisone (Sigma-Aldrich), 5 ug/mL Insulin (Gibco-), 8.4 ng/mL Cholera toxin (Sigma-Aldrich), 10 ng/mL Epidermal growth factor (BPS bioscience), 10 umol/L Y-27632 (Abmole), 50 ug/mL Gentamicin (Gibco), 1% P/S (Thermo Fisher Scientific,), Amphotericin B (1 ug/ml) (Thermo Fisher Scientific). After five passages of coculture, murine irradiated 3T3-J2 cells were removed using a Feeder Removal MicroBeads kit (Miltenyi), and epithelial cancer cells were expended in F-media. PDX-derived cell lines were cultured in F media: 5% fetal bovine serum, 400 ng/ml hydrocortisone, 5 μg/ml insulin, 8.4 ng/ml cholera toxin, 10 ng/ml hEGF, 10 μM Y-27632 (ROCK inhibitor), 50 μg/ml gentamicin.

All cell lines used were routinely tested for Mycoplasma (Lonza Mycoalert and EZ-PCR Mycoplasma Detection Kit) and were authenticated using short tandem repeat analysis. The human origin of PDX-derived cell lines was validated by flow cytometry using FITC anti-human EpCAM antibody clone VU-1D9 and (Thermo Fisher Scientific, #A15755) and PE/Cy7 anti-mouse H2Kd antibody clone SF1-1.1 (Biolegend, # 116622). All cells were maintained at 37°C, 5% CO_2_.

### Single and dual-gene overexpression assay

For generation of stable *c4orf19, RBPJ, SEPSECS, SEL1L3, KCNIP4, TBC1D19, RELL1, SLC34A2, STIM2, LGI2, PI4K2B* and *CCKAR* overexpression cells, lentiviral ORF vectors were retrieved from the arrayed MGC premier human lentiviral ORF (Sigma) (ccsb ID, blasticidin resistant) and MISSION® TRC3 Human ORF collection (Sigma) (TRCN ID, puromycin resistant) obtained from the McGill Platform for Cell Perturbation (MPCP). The following lentiviral ORF vectors were used: *c4orf19* (ccsbBroad304_03572, TRCN0000469204), *RBPJ* (ccsbBroad304_06435, TRCN0000470066), *SEPSECS* (ccsbBroad304_11945), *SEL1L3* (ccsbBroad304_11701, TRCN0000479888), *KCNIP4* (ccsbBroad304_09030, TRCN0000474912), *TBC1D19* (TRCN0000468467), *RELL1* (TRCN0000476648), *SLC34A2* (TRCN0000476745), *STIM2* (TRCN0000477969), *LGI2* (TRCN0000481617), *PI4K2B* (TRCN0000489163), *CCKAR* (TRCN0000489014, TRCN0000491970), *CDKN1A* (ccsbBroad304_00282, TRCN0000471863), *CDKN1B* (ccsbBroad304_05980, TRCN0000475049) and GFP control vector pLX317-GFP and pLX304-GFP. Viral particles were produced by co-expressing ORF or control constructs with packaging plasmids psPAX2 and pMD2.G in HEK-293T cells using lipofectamine 2000 transfection protocol. Media containing viral particles was collected and passed through a 0.45 μm filter. Cells were treated with virus in media containing 8 μg/ml polybrene. Twenty-four hours after transduction cells were recovered for another 24 hrs and then MCF10A were selected in 3 μg/ml puromycin dihydrochloride (Sigma) for 48 hrs or 10 μg/ml blasticidin (Gibco) for 72 hrs for; GCRC1735 in 5 μg/ml puromycin dihydrochloride for 48 hrs or 7.5 μg/ml blasticidin for 72 hrs and GCRC1915 in 5 μg/ml puromycin dihydrochloride for 48 hrs or 10 μg/ml blasticidin for 72 hrs. MOI was determined for each construct for each cell line and ∼ 0.3 MOI was used for all constructs ensuring one integrant per cell. Viral transductions with the respective vectors were carried out sequentially. Overexpression was confirmed by Western Blot using Anti-V5 antibody (Abcam #27671, 1:1000). All single gene overexpression mutant cells were constructed such that they overexpressed each ORF under one of the selection markers and GFP was then under the second selection marker. All double gene overexpression mutant cells were constructed such that they overexpressed each ORF under one of the selection markers and the second ORF was then under the second selection marker. Confluence was measured by IncuCyte. Briefly, 4000 MCF10A and GCRC1915 cells and 2000 GCRC1735 cells were plated per well representing 20% confluence at the start of the experiment. Imaging was done at 4 hr intervals for a duration of 5-6 days. Each mutant cell line was plated in three wells per plate for a total of three technical replicates. The experiment was repeated for a total of three independent biological replicates. *CDKN1A* and *CDKN1B* were included as positive controls, which are known to lead to a severe proliferation defect when overexpressed^19^.

### Genome-wide pooled overexpression screen

The pooled MISSION TRC3 LentiORF collection (Sigma) provided by the McGill Platform for Cell Perturbation (MPCP) was used to infect MCF10A (10^8^) and GCRC1735 (1.5×10^8^) cell lines. Cells were treated with virus in media containing 8 μg/ml polybrene for 24 hrs. Viral supernatant was removed and the media was refreshed recovering the cells for 48 hrs and then MCF10A and GCRC1735 were selected in 3 or 5 μg/ml puromycin dihydrochloride for 48 hrs, respectively. MOI ∼ 0.3 MOI was used for the screens and 1000x coverage was maintained at each step of the screen for both cell lines. Following 6-8 doublings, genomic DNA was isolated using the Roche High Pure PCR Template Preparation Kit followed by an RNase A treatment. One microgram of DNA was then used in 48 2-step PCR reactions with barcoded Illumina sequencing primers and then with P5/P7 primers. The reactions were then purified using the Roche PCR Purification Kit. Samples were then sequenced at The Center for Applied Genomics at Toronto Sick-Kids hospital on the Illumina HiSeq 2500 platform. The 50-base kit with 62 cycles and single-end reads was used to obtain the exact read-length needed for the library vector. Sequences were then deconvoluted. For all downstream analyses, we only included genes with a read count higher than 100 in T0 samples (MCF10A_T0 and GCRC1735_T0). Raw counts were normalized using edgeR’s TMM algorithm (Robinson et al., 2010) and were then transformed to log2-counts per million (logCPM) using the voom function implemented in the limma R package (Ritchie et al., 2015). To assess differences in gene expression levels, we fitted a linear model using limma’s lmfit function. Nominal p-values were corrected for multiple testing using the Benjamini-Hochberg method. Genomic heatmaps of log2 fold-changes were created using CNVkit (Talevich et al., 2016).

### TCGA data computational analysis

Patient copy number data were obtained from a TCGA Breast Invasive Carcinoma (BRCA) (n = 2199) using Firehose Broad GDAC (https://gdac.broadinstitute.org/; accessed on 31 July 2016). Frequencies of gene deletions were derived from the single nucleotide polymorphism array dataset (genome_wide_snp_6-segmented_scna_minus_germline_cnv_hg19) and analyzed by GISTIC2.0 (Mermel et al., 2011). Parameters used for analysis were: reference genome build hg19; amplification threshold 0.1; deletion threshold -0.1; join segment size 4; qv threshold 0.25; remove X chromosome yes; cap value 1.5; confidence level 95; broad analysis yes; broad length cut-off 0.5; maximum samples per segments per sample 2000; arm peel-off yes. PAM50 annotation was obtained from a previous study ^8^. TP53 mutations were obtained from cBioPortal and missense mutations were annotated by IARC filename: functionalAssessmentIARC TP53 Database, R18.xlsx. LOF mutations were considered: Frame_Shift_Del, Frame_Shift_Ins, Nonsense_Mutation, Splice_Site and Missense_Mutation if there were more cases of LOF than GOF.

Patient copy number data were obtained from a TCGA Lung Squamous Cell Carcinoma (LUSC, n = 1032), Testicular Germ Cell Tumors (TGCT, n = 304), Ovarian Serous Carcinoma (OV, n = 1168), Esophageal carcinoma (ESCA, n = 373), Cervical Squamous Cell Carcinoma and Endocervical Adenocarcinoma (CESC, n = 586), Mesothelioma (MESO, n = 172), Rectum adenocarcinoma (READ, n = 316), Stomach Adenocarcinoma (STAD, n = 904), Breast Invasive Carcinoma (BRCA, n = 2199), Colon Adenocarcinoma and Rectum Adenocarcinoma (COADREAD, n = 1234), Colon Adenocarcinoma (COAD, n = 918), Head and Neck Squamous Cell Carcinoma (HNSC, n = 1089) and Uterine Corpus Endometrial Carcinoma (UCEC, n = 1089) (https://gdac.broadinstitute.org/; accessed on 31 July 2016). Frequencies of gene deletions were derived from the single nucleotide polymorphism array dataset (genome_wide_snp_6- segmented_scna_minus_germline_cnv_hg19) and analyzed by GISTIC2.0 (Mermel et al., 2011). Parameters used for analysis were: reference genome build hg19; amplification threshold 0.1; deletion threshold -0.1; join segment size 4; qv threshold 0.25; remove X chromosome yes; cap value 1.5; confidence level 95; broad analysis yes; broad length cut-off 0.5; maximum samples per segments per sample 2000; arm peel-off yes. Higher amplification and deletion thresholds than above were used to increase stringency for the pan-cancer analysis.

TCGA gene expression data set from breast cancer invasive ductal carcinoma was used for differential gene expression. For all downstream analyses, excluded lowly expressed genes with an average read count lower than 10 were excluded from all samples. Raw counts were normalized using edgeR’s TMM algorithm and were then transformed to log2-counts per million (logCPM) using the voom function implemented in the limma R package. To assess differences in gene expression levels, we fitted a linear model using limma’s lmfit function. Nominal p-values were corrected for multiple testing using the Benjamini-Hochberg method. Gene-Set enrichment analysis based on pre-ranked gene list was performed using the R package fgsea (http://bioconductor.org/packages/fgsea/). Default parameters were used.

### Immunoprecipitation and Western Blot

MCF10A cells were plate at a density of 1.5×10^6^/10cm-dish and transfected with 4µg of candidate interactors using Lipofectamine (ThermoFisher, 18324012) and Plus (ThermoFisher, 11514015) transfection reagents. For this DNA constructs were incubated with 8µl Plus reagent in 500µl Opti-MEM (ThermoFisher,11058-021), Lipofectamine reagent was incubated separately in another 500µl Opti-MEM, for 15min; following initial incubation, the solutions were mixed and incubated together for another 15min. The transfection mixture was added dropwise to pre-washed cells containing 2ml Opti-MEM, and incubated for another 3h at 37˚C, at which point the transfection solution was removed, and cells were returned to normal growth media. Following 24hrs, cells were harvested in lysis buffer (50 mm HEPES, 150 mm NaCl, 1.5 mm MgCl_2_, 1 mm EGTA, 1% Triton X-100, 10% glycerol 1 mM PMSF, 1 mM Na_3_VO_4_, 1 mM NaF, 10 μg/ml aprotinin and 10 μg/ml leupeptin, pH 7.4). Lysates were pre-cleared with 30µl of either protein-A-sepharose (GE Healthcare, 17-5280-01) or protein-G-sepharose beads (GE Healthcare, 17-0618-01) for 1h at 4˚C. 1500µg of protein was then incubated with either 1.8μl (*∼5µg*) V5-tag antibody (Abcam, ab27671), 1.25µl (*∼5µg*) FLAG-tag antibody (Sigma, F3165) or 1.5µl (*∼5µg*) mouse-IgG negative control and either 40µl of protein-A or protein-G-sepharose beads overnight at 4°C. Beads with bound proteins were washed three times in lysis buffer plus inhibitors, and eluted by boiling in SDS sample buffer. Eluted proteins and 50 μg of protein from whole cell lysate were resolved in 4-15% NuPAGE gradient gel (ThermoFisher, NP0335) using MOPS running buffer (ThermoFisher, NP000102). Proteins were transferred on PVDF Odyssey membranes (MilliporeSigma) using a Mini Trans-Blot System from Bio-Rad. Detection and quantification of protein levels was performed on the Odyssey IR imaging System (Li-COR Biosciences) using fluorescently labeled secondary antibodies, anti-mouse-680 (Mandel Scientific, LIC-926-68070) or anti-rabbit-800 (Mandel Scientific, LIC-926-32211).

### miniTurboID

Gateway cloning was used to clone c4orf19 (ccsbBroadEn_03572) from pDONR223 to pSTV6- miniTurbo. MCF10A cells were transduced with lentivirus backbone containing pSTV6- C4orf19-3xFLAG-miniTurbo or pSTV6-GFP-3xFLAG-miniTurbo and selected in media containing puromycin as previously described in the “Cell Culture” section. Western blot was conducted to ensure C4orf19-3xFLAG-miniTurbo expression and biotinylation as described above. Anti-GAPDH primary antibody (Santa Cruz Biotechnology #sc-25778; 1:1000) was used as described above. Streptavidin-HRP conjugate (Millipore Sigma #RPN1231VS, 1:5000) was used and visualized using Immobilon Forte Western HRP substrate (Millipore Sigma #WBLUF0500).

Cells were grown to ∼70% confluency and bait expression and biotin labeling was induced simultaneously (0.5 μg/ml doxycycline, 40 μM biotin). After 4 h, cells were rinsed and scraped into 1 mL of PBS. Cells were collected by centrifugation (500 × g for 3 min) and stored at −80 °C until further processing.

Cell pellets were thawed on ice and resuspended in lysis buffer containing 50 mM Tris-HCl, pH 7.5, 150 mM NaCl, 1% Nonidet P-40 substitute (NP40; IGEPAL-630), 0.4% SDS, 1 mM MgCl_2_, 1 mM EGTA, 0.5 mM EDTA, 0.4 % sodium deoxycholate, benzonase & protease inhibitors at a ratio of 10:1 (w/v). Cells were lysed with 15 seconds of sonication (5 sec on, 3 sec off) at 30% amplitude on a Q500 Sonicator with an 1/8” Microtip and were rotated end-over-end at 4 °C for 20 min. Cell debris was pelleted via centrifugation at 15,000 × g for 15 min at 4 °C. Supernatants were incubated with 25 μL (packed bead volume) of streptavidin-Sepharose beads (GE) with rotation for 3 hr at 4 °C. Beads were pelleted at 500 × g for 2 min, transferred to new tubes and resuspended in 500 μL of fresh lysis buffer.

Beads were washed once with SDS wash buffer (50 mM Tris-HCl, pH 7.5, 2% SDS), twice with lysis buffer, once with TNNE wash buffer (50 mM Tris-HCl, pH 7.5, 150 mM NaCl, 1 mM EDTA, 0.1 % NP-40), and thrice with 50 mM ammonium bicarbonate, pH 8.0 (ABC). Each wash consisted of bead resuspension in 500 µl of each buffer, pelleting of beads at 500 × g for 30 sec and aspiration of supernatant. On-bead digestion was performed by resuspending beads in 100 μL of ABC containing 1 µg of sequencing grade trypsin (T6567, Sigma-Aldrich). Samples were gently mixed at 37 °C overnight. Samples were spiked with 1 µg of fresh trypsin and digested further for 3 hr. The supernatant, containing digested peptides, was transferred to new tubes. Beads were washed twice with HPLC-grade water to wash off peptides, and these were pooled with the collected supernatant. Peptides were vacuum centrifuged until dry.

#### Mass spectrometry acquisition

Each sample was resuspended in 5 % formic acid and loaded onto an equilibrated high-performance liquid chromatography column (800 nL/min). Peptides were eluted with a 90 min gradient generated by a Eksigent ekspert™ nanoLC 425 (Eksigent, Dublin CA) nano-pump and analyzed on a TripleTOF™ 6600 instrument (AB SCIEX, Ontario, Canada).

The MS acquisition method has been described previously on identical instrumentation ^70^. The gradient was delivered at 400 nL/min and consisted of three steps: sample delivery, column cleanup and column equilibration. The gradient used to pass sample over the column took place over 90 min starting with 2% acetonitrile (ACN) + 0.1 % formic acid (FA) and ending with 35 % ACN + 0.1 % FA. Cleanup was performed by passing 80 % ACN + 0.1 % FA over the column for 15 min, and the column was equilibrated back to 2 % ACN + 0.1 % FA over 15 min.

Instrument calibration was performed on bovine serum albumin reference ions to adjust for mass drift and verify peak intensity before samples were analyzed in data-dependent acquisition (DDA) mode. One 250 ms MS1 TOF (time of flight) survey scan (over mass range 400 - 1800 Da) was performed and was followed by 10 × 100 ms MS2 candidate ion scans (100 - 1800 Da). Ions that exceeded a threshold of 300 counts per second and had a charge of 2+ to 5+ were selected for MS2. Precursors were excluded for 7 sec after one occurrence.

#### Data-dependent acquisition data search

The ProHits laboratory information management system was used to analyze proteomics data ^71^. WIFF files were converted with the WIFF2MGF converter and to a mzML format using ProteoWizard (V3.0.10702) and the AB SCIEX MS data converter (V1.3 beta). Converted files were searched with Mascot (2.3.02) ^72^ & Comet (2016.01 rev.2) ^73^. Spectra were searched against a collection of 72,482 entries comprised the following: human and adenovirus sequences (version 57, January 30th, 2013), common contaminants [Max Planck Institute (http://maxquant.org/contaminants.zip) & Global Proteome Machine (GPM; ftp://ftp.thegpm.org/fasta/cRAP/crap.fasta)], reversed sequences, bait tags (eg. BirA or GFP) and streptavidin. Search parameters were set to search for trypsinized peptides allowing for two missed cleavages. For precursors, a mass tolerance of 35 parts per million was set, and peptides of +2 to +4 charges were allowed with a tolerance of ± 0.15 amu for fragment ions. Variable modifications included deamidated asparagine and glutamine as well as oxidized methionine. Search results were analyzed with the Trans-Proteomic Pipeline (v4.7 POLAR VORTEX rev 1) and iProphet pipeline ^74^.

#### SAINT analysis

An iProphet probability score > 0.95 and more than two unique peptides were required for protein identification. SAINTexpress [version 3.6.1 ^75^] was used to score proximity interactions from DDA data using default parameters. Bait runs, run in biological duplicate, were compared against four negative control runs consisting of two miniTurbo-eGFP-only samples and two untransduced MCF10A samples. Control runs were not compressed for this analysis. Preys with a Bayesian false discovery rate < 5% were considered high-confidence proximity interactions. gProfiler (https://biit.cs.ut.ee/gprofiler/gost) was used to calculate enrichment of GO cellular component terms.

### RNA *in situ* hybridization and immunofluorescence of GCRC1735

FFPE tissue was deparaffinized and underwent heat-mediated antigen retrieval in citrate buffer pH6.0 or EDTA buffer pH9.0. Slides were blocked with Power Block for 5 min at room temperature, and incubated with the primary antibody for 30 min at room temperature followed by washing with TBST (3 x 3min). Slides were incubated with secondary antibody-HRP for 30 min at room temperature, washing with TBST 3x 3min and stained with Opal fluorophore working solution for 10 min. This was followed by heat-mediated antibodies stripping to remove the primary and secondary antibodies in order to repeat additional rounds for labeling with other primary antibodies. The primary antibodies are against Ki67 (Ventana #790-2910) and Pan-Keratin (Cat# 760-2595, Ventana). The antibody specificity and dilution were tested before multiplex assay. Nuclei were stained with 0.5 ng/ml DAPI for 5 minutes at room temperature and counterstaining was done with Harris’ hematoxylin. RNA *in situ* hybridization was performed using the RNAscope 2.5 HD Assay (cat#322360. ACD Bio) according to the manufacturer’s instruction on FFPE PDX section. The probes used are Hs-RBPJ (cat#448661), the positive control Hs-PPIB housekeeping gene and the negative control dapB. Slides were imaged with an LSM800 confocal microscope (Zeiss). Brightfield slides were scanned using Aperio-XT slide scanner (Aperio). Visual inspection was used to classify cells into RBPJ or Ki67 high and low classes. Due to low number of RBPJ high expression cells a field with equal number of cells of both RBPJ expression classes was used for the quantification.

### C4ORF19 immunofluorescence and quantification

C4orf19 immunofluorescence staining: cells were seeded in 24-well plate with coverslips until they reached 80-90% confluence. Then, they were fixed in 4% paraformaldehyde (20 min), permeabilized with 0.2% Triton X-100 (10 min), blocked with 2% BSA (30 min), and then incubated with Anti-C4orf19 primary antibody (1:100, GeneTex, GTX106538) (1 hr). The primary antibody was visualized with a fluorescent secondary antibody conjugated to Alexa Fluor 488 raised in goat (1:1000, Invitrogen A21206) (1 hr). Nuclei were counterstained with 0.25 ng/ml DAPI (5 min). All steps were performed at room temperature. Images were acquired on the Nikon C2/TIRF confocal laser scanning microscope (Nikon), using a 63X objective.

Fiji (v.2.3, NIH) was used to analyze subcellular localization of C4ORF19 using a custom macro. The DIC image was used to manually outline each cell; DAPI was used to create a nuclear mask and GFP channel was used to quantify C4ORF19 subcellular localization. The macro makes bands of ∼3µm from the edge of a cell outline into the middle of the cell, and measures the mean intensity, total amount of signal, the proportion of the cell’s total area that is in this band, the proportion of the cell’s total signal that is in this band, and the ratio of the signal-to-area. The ratio of the signal-to-area is above 1, if there is a greater proportion of the signal in that band than might be expected based solely upon area. The outer band representing ∼ 25% of the cell area was compared to the remainder of the cell to quantify the protein abundance of C4ORF19 in the cell periphery compared to the cytoplasm.

### Single cell RNA sequencing

Breast mammoplasty reduction epithelial single-cell suspensions were washed three times in PBS with 0.04% BSA. An aliquot of cells was used for LIVE/DEAD viability testing (Thermo Fisher Scientific). Single-cell libraries were generated using the Chromium Controller and Single Cell 3’ Library & Gel Bead Kit v3 and Chip Kit (10x Genomics) according to the manufacturer’s protocol. Briefly, cells suspended in reverse transcription reagents, along with gel beads, were segregated into aqueous nanoliter-scale gel bead-in-emulsions (GEMs). The GEMs were then reverse transcribed in a T1000 Thermal cycler (Bio-Rad) programed at 53°C for 45 min, 85°C for 5 min, and hold at 4°C. After reverse transcription, single-cell droplets were broken and the single-strand cDNA was isolated and cleaned with Cleanup Mix containing DynaBeads (Thermo Fisher Scientific). cDNA was then amplified with a T1000 Thermal cycler programed at 98°C for 3 min, 12 cycles of (98°C for 15 s, 63°C for 20 s, 72°C for 1 min), 72°C for 1 min, and hold at 4°C. Subsequently, the amplified cDNA was fragmented, end-repaired, A-tailed and index adaptor ligated, with SPRIselect Reagent Kit (Beckman Coulter) with cleanup in between steps. Post-ligation product was amplified with a T1000 Thermal cycler programed at 98°C for 45 s, 12 cycles of (98°C for 20 s, 54°C for 30 s, 72°C for 20 s), 72°C for 1 min, and hold at 4°C. The sequencing-ready library was cleaned up with SPRIselect and quantified by qPCR (KAPA Biosystems Library Quantification Kit for Illumina platforms). 200 pM of sequencing libraries were loaded on an Illumina HiSeq instruments (see Single-cell RNA sequencing analysis section) and ran using the following parameter: 26 bp Read1, 8 bp I7 Index, 0 bp I5 Index and 98 bp Read2.

### Single-cell RNA sequencing analysis

Two samples were sequenced using the Chromium single cell 3’ RNA-seq on 0.5 lane the NovaSeq S1 instrument, for a total of 492,626,914 reads, and 327,762 reads per single-cell (saturation 89.2%). Alignment against the human GRCh38 genome was performed using Cell Ranger Pipeline version 3.0.1. The Ensembl annotation for GRCh38 (release 93) was used, keeping only the genes with the biotypes protein_coding, lincRNA and antisense. Empty GEMs containing only background reads were discarded by the pipeline and bar code errors resulting from sequencing were corrected if they contained only one mismatch by assigning them to the closest available bar code, or discarded otherwise, resulting in 1,503 GEMs containing cells. Alignment quality was controlled by assessing the proportion of reads mapping confidently to the transcriptome (52.4%). The total number of genes detected (>1 mapped read) was 20,541, with a median of 880 per cell. The R package Seurat (v3.2.3) was used to analyze the single-cell RNA-seq data (Satija et al., 2015). Cells with over 12% mitochondrial content, over 40,000 UMIs, or less than 500 UMIs were discarded. Gene counts were normalized to a total of 10,000 UMIs for each cell, and transformed to a natural log scale. Counts were then adjusted for library size and mitochondrial proportions. Heat Digestion Stress Response Gene Set as previosuly reported ^76^ was identified and removed. Cell types were annotated using previously defined markers ^76^ (Fig. 4S). Cells in the breast epithelial cell cluster were used as a baseline of normal gene expression for inferring copy number described below.

### Inferring copy number aberrations from scRNAseq

Copy number profile was inferred from scRNAseq data as previously described ^33^. Briefly, the gene expression was normalized to ensure cells are comparable, whereby the Trimmed-Mean M normalization rescale expression in a cell by a factor to match that of a control cell. Consecutive genes were merged to form “bins” with a minimum average expression. Each bin was normalized across cells to produce a Z-score which was computed by subtracting the average expression across all the cells and dividing by the standard deviation. The normalized expression was smoothed using a rolling median approach: the expression in each bin was replaced by the median expression of the surrounding bins in the cell. Specifically, the rolling median was run-in windows of size 5, i.e., the bin of interest and two bins on each side. The smoothed Z-score was winsorized to be within [-3,3] to further reduce the effect of single-gene outliers. After this step, the score is centered on 0 and positive (negative) values support a higher (lower) copy number than in the majority of the cells. A principal component analysis (PCA) was run on the smoothed Z-scores. To minimize the effect of cell cycle, the PCA was run on non-cycling cells and all the cells projected on this principal components (PCs).

Community detection was performed by the Louvain algorithm on a cell network built from the top PCs. A tSNE was run on the top 20 PCs. To build the cell network, we first identify the K-nearest neighbors of each cell based on the Euclidean distance D in the PC space. The K-nearest neighbor cells are linked in the network with a weight defined as 1/(1+D). The Louvain community detection was then run on this cell network. Gamma resolution parameter with a high mean Rand Index across the runs and/or a low Rand Index variance was used. Copy-number aberrations are called at the community level to increase the sensitivity to shorter aberrations. Meta-cells were constructed by combining the expression of randomly selected cells in a community. We created multiple meta-cells for each community and looked for consistent CNA signal in all meta-cells. The expression in each meta-cell was normalized similarly as for the CNA-based community detection: normalization per cell, merging into expressed bins, Z-score computation, and smoothing. The Z-score was computed relative to a specific baseline, e.g., cells identified as normal that were isolated from mammoplasty reduction. CNA were called using an HMM with three states: neutral copy number, loss, and gain. A Gaussian mixture HMM was capable of segmenting together the multiple meta-cells from a community. Short copy-number segments are filtered, for example if spanning less than 5 consecutive bins, as they could result from single genes with strong expression differences. A Wilcoxon test was performed to assess the significance of each loss/gain segment by comparing the expression within the segment with the expression in nearby “neutral” segments.

### WGS analysis

Bulk WGS data for PT and PDX including BAM generation, Manta calls for structural variants and Mutect2 calls for somatic mutation variants for were obtained from a previous study_29._

### Timing analysis using bulk WGS data for basal breast cancer PT/PDX panel

#### CNA profiles

CNA profiles were obtained using Battenberg (v2.2.9) ^77^, integrating SV calls from Manta and correcting logR for both GC content and replication timing. SNP phasing was performed using Beagle 5.1 (18May20.d20). To better capture LOH-related events in PDX samples, because BAF in pure samples tends to be highly squished and might be misinterpreted, purity was artificially decreased by pulling allele counts from both germline and PDX and was set back to 100% when checking sample purity with SNV information from Mutect2. All CNA profiles were manually examined and quality checked (*e.g.* homozygous deletions, superclonal peaks, purity estimates, etc.). Whole-genome doubling (WGD) information was assessed using the relationship between the fraction of the genome with LOH and ploidy, as in PCAWG studies ^78^.

#### CNA clustering

Clustering of CNA profiles was performed using MEDICC2 (v0.3) ^79^. Genomic regions of more than 500kb and covered in all samples for a given patient were considered, with *cn_a* and *cn_b* defined as the major and the minor allele, respectively. The copy-number state of the most abundant subclone was selected for subclonal CNAs. Since WGDs were clonal events, we defined reference normals with a 1+1 baseline in samples without WGD and 2+2 with WGD so the output was WGD-aware.

#### Subclone trees

Subclonal compositions were assessed using DPClust (v2.2.8) ^77^ and SNVs information from Mutect2 calls, leveraging principles of reconstructing subclone trees ^80^. Small and noisy clusters (<5% of total SNVs) breaking the pigeonhole principle were discarded from the final trees.

#### Genomic Event Timing

Clonal copy number gains were timed using an approach similar to that outlined in a previous study ^17^. A posterior distribution over SNV multiplicities was measured using the emcee sampler (https://arxiv.org/abs/1202.3665) with a prior that corresponded to a uniform distribution over gain timing. For each segment we ran 30 independent chains for 2000 steps with 1000 burnin steps. The posterior distribution over SNV multiplicities was converted into a distribution over gain timing.

The timing of the WGD for WGD tumors was measured by jointly timing all the gains that resulted in a major copy number state of two. For WGD samples, the timing of gains leading to major copy number three and four states were measured with equal prior probability on a gain occurring before or after the WGD. The relative likelihood of pre-or post-WGD gains was calculated by measuring the similarity between the segment WGD timing distribution measured with the route compared to the sample-wide WGD timing distribution. For gain regions with major copy number four, the timing of the average of the two post-WGD gains was measured as the system is underdetermined. Only gained regions with a major copy number of up to four in WGD tumors and two in non-WGD tumors were timed. The gained regions also needed to have at least 10 SNVs and a minor copy number of no more than two. The timing of SNVs in key genes was measured using MutationTimeR ^17^.

The PDXs were used to refine our timing of events in the primary tumor. If an event was identified as clonal in the primary tumor, but was not found in the PDX, we reclassified the event as subclonal in the primary as the event was likely not present in the cells from the primary that seeded the PDX.

#### League Model

A league representing a timeline of genomic events aggregated across tumors were produced from our timing data using a league modeling approach similar to that outlined in previous studies^17,81^. Briefly, the aggregate timing of the events is determined by running a scoring process where the earliest events accumulate the higher score. This is achieved by initialising each genomic event with a score of zero. We then sample the relative timing for each possible pair of genomic events from the subset of individual tumor timelines in our cohort that contain both events. The earlier event has its score increased by one and the later event decreased by one. If the relative timing of the event pair cannot be distinguished, or if no sample has both events, the score for both events is kept the same. After each possible pair of events is considered, the events are ranked according to their score. This process is repeated 100 times to achieve a distribution over the ranks.

#### Real-Time Timing

A real-time estimate of WGD and the emergence of the MRCA was achieved using the approach outlined in a previous study ^17^. Instead of evaluating the timing using both a branching and linear subclonal structure ^17^ we used the structure inferred from our subclone tree reconstruction.

### scDNAseq

Basal breast cancer PDX-derived GCRC1735 single-cell suspensions were washed three times in PBS with 0.04% BSA. An aliquot of cells was used for LIVE/DEAD viability testing (Thermo Fisher Scientific). Single-cell DNA libraries were generated using the Chromium Single Cell DNA Reagent Kit (10X Genomics) according to the manufacturer’s protocol. Briefly, an appropriate volume of cell suspension for targeting 500 cells were added to the Single Cell Bead Mix then loaded onto a Chromium Chip C, along with CB polymer. The resultant Cell-Bead was allowed to polymerize overnight shaken at 1000rpm. The encapsulated cells were then lysed and its genomic DNA was NaOH denatured. The Cell-Bead along with a reaction mix and Gel-Bead were loaded onto the Chromium D Chip to generate gel bead-in-emulsions (GEMs) on the Chromium Controller. An ideal GEM will contain reaction mix, one Cell-Bead and one Gel-Bead. The GEMs were incubated in a T1000 Thermal cycler (Bio-Rad) programed at 30°C for 3hour, 16°C for 5 hour, 65°C for 10 min, and hold at 4°C. Then the GEMs were broken and its amplified DNA were isloated using Dynabeads MyOne Silane beads followed by a SPRIselect cleanup. DNA was quantified on a Caliper Labchip (Beckman Coulter) using High Sensitivity DNA Assays. The DNA was converted to sequence ready library by fragmentation, end-repaired, A-tailing, index adaptor ligated and index PCR with SPRIselect clean ups in between. Four samples were sequenced on 2 lanes the Illumina HiSeqX instrument, for a total of 3,726,525,082 reads, and 505,266 reads per single-cell (saturation 15%).

### scDNAseq data processing

Sequencing data was processed by using 10X Cell Ranger DNA pipeline to generate a raw bam for each sample. Briefly, the reads were aligned to the human reference genome build 38 (GRCh38) by using BWA and then converted to sorted BAM. The bam file was demultiplexed into individual bam files by using in-house python script to represent the sequencing reads from each single cell. Poorly mapped reads with mapping quality < 25 were filtered out by using SAMtools. PCR duplicates were removed by using Picard. Noisy cells detected by 10X Cell Ranger DNA pipeline with depth independent MAPD statistically higher than the sample distribution (with p-value < 0.01) and low ploidy confidence were excluded. We also filtered out cell outliers with large Lorenz curve area ^82^.

### scDNAseq SNV analysis

To create a pseudo-bulk sample, single-cell DNA (scDNA) samples were merged while preserving the cell of origin information. This was achieved by incorporating the cell of origin information into the read group field of each read. The pseudo-bulk sample was then processed as a standard whole-genome sequencing (WGS) sample using the tumor_pair pipeline from Genpipes ^83^. Somatic variants were generated using Mutect 2 and were utilized for single-nucleotide variant (SNV) fishing in individual cells. To reduce the false positive rate during the fishing process, we excluded indels and retained only high-quality somatic variants (TLOD >= 40). For each cell (represented by each read-group in the pseudo-bulk), we extracted the base distribution at each selected somatic position using BVAtools basefreq (https://bitbucket.org/mugqic/bvatools/src/master/). Cell-specific somatic variants were determined by comparing the extracted base frequency with the expected variant allele detected by Mutect 2. In order to ensure robustness, cell-specific somatic variants were excluded if they were not genotyped in at least 10 different cells.

### Building the clone tree and assigning SNVs to clones using heuristics

SNVs were used to build the clone tree for GCRC1735 primary and PDX tumors. SNVs were independently called in four datasets: primary tumour bulk (PT_bulk), PDX bulk (PDX_bulk), single-cell samples 1 and 2 pseudobulk (SCS12) and single-cell samples 3 and 4 pseudobulk (SCS34). After examining the patterns of SNVs presence and absence in the samples, and based on the known ancestral relationships among the sample, the following heuristics were developed to assign mutations to clones (**Fig S3A**). First, if a mutation was present in all datasets, it was deemed a clonal mutation. Second, if a mutation was present in PT_bulk and SCS12 and not present in SCS34 or PDX_bulk, then it was assigned to subclone 1. Third, if a mutation is present in PT_bulk, PDX_bulk and SCS34 but not in SCS12, then it is set to subclone 2. Fourth, if a mutation is present only in PDX_bulk and SCS34 then it is set to subclone 3. Due to the low count of unique mutations to PT_bulk or PDX_bulk, we did not attempt to further define smaller subclones present in these samples.

### Inference of haplotype-specific copy number profiles in single cells using CHISEL

CHISEL^84^ is a tool that infers haplotype-specific copy number profiles of each cell in a low coverage single-cell sequencing dataset. This was used on the single-cell datasets to determine copy number profiles for subclones 1 and 3. To run CHISEL we first installed the package and dependencies as instructed on their GitHub page. Germline SNPs were called using bcftools’ `mpileup` and `call` methods on the normal sample .baf file. These SNPs were phased using the Michigan Imputation Server (MIS). As required for MIS input, the SNP calls were separated by chromosome into separate .bcf files, sorted by genomic position, and uploaded to their server. For phasing on MIS, we used as arguments: reference panel `HRC r1.1 2016 (GRCh37/hg19)`, array build `GRCh38/hg38`, phasing `Eagle v2.4`, and mode `Quality control and phasing only`. The MIS results were downloaded as a set of chromosome-specific .vcf files and merged. The X chromosome was omitted during this merge as, at the time of running, sex chromosomes were not permitted in CHISEL’s input. The MIS outputs phased SNP data using hg19 reference genome coordinates. Therefore, these coordinates were lifted over to the hg38 reference genome. This was done using picardtool’s `LiftoverVcf` function with the requisite UCSF chain file. CHISEL also requires a .bam file for the single-cell sequencing data. In CHISEL, haplotypes are categorized as maternal or paternal arbitrarily, and so single-cell datasets could not be input to CHISEL separately. Instead, all single-cell data files were merged into a single .bam file and used as input. With these phased germline SNPs and the single-cell .bam file, CHISEL was run with default settings. This process was repeated using a different seed to confirm reproducibility.

### Inferring the Most Recent Common Ancestor (MRCA) copy number profile

The results from CHISEL reveal two predominant subclones, each defined by the PDX line the cells are derived from. To estimate the haplotype-specific copy number profile of their MRCA, we developed and applied the following heuristics. First, if an allele’s copy number was the same between the two subclones, that allele’s copy number was set to the same for the MRCA. Second, if there was a LOH event in one subclone but not the other, the MRCA copy number state was set to contain the lost allele. Third, in other regions of differing copy number, if there was an adjacent region with shared copy number between the two subclones, and that region’s copy number matched that of one of the two mismatched copy numbers, then the MRCA copy number for the differing region was set to be the same as the matching neighbour. If none of the previous three heuristics apply, then the MRCA copy number is set to be the minimum of the two subclonal copy numbers.

## Supplementary tables

**Table S1.** GISTIC2.0 Broad deletion analysis in TCGA basal breast cancer cohort.

**Table S2.** Differential gene expression of chr4p genes in TCGA basal breast cancer cohort with deletion or copy neutral status of chr4p.

**Table S3.** GISTIC2.0 Broad deletion analysis in TCGA pancancer cohort.

**Table S4.** Transcriptome-wide differential gene expression in TCGA basal breast cancer cohort with deletion or copy neutral status of chr4p.

**Table S5.** Aneuploidy in basal breast cancer with different chr4p copy number states. Aneuploidy score as quantified by Chrom.Arm.SCNA.Level median reported by Davoli et al Science 2017.

**Table S6.** Phylogenetic reconstruction using bulk WGS data of PT/PDX basal breast cancer cohort.

**Table S7.** Phylogenetic reconstruction using scDNAseq data of PT/PDX GCRC1735 basal breast cancer.

**Table S8.** Gene expression and cell clusters as identified from scRNAseq of normal breast epithelial tissue.

**Table S9.** Relationship between transcriptional programs and inferred copy number changes using scRNAseq data of GCRC1735 PDX.

**Table S10.** miniTurboID screen for C4orf19.

**Table S11.** Gene overexpression screen for MCF10A and GCRC1335 PDX-derived cell line.

---

**Figure S1.**
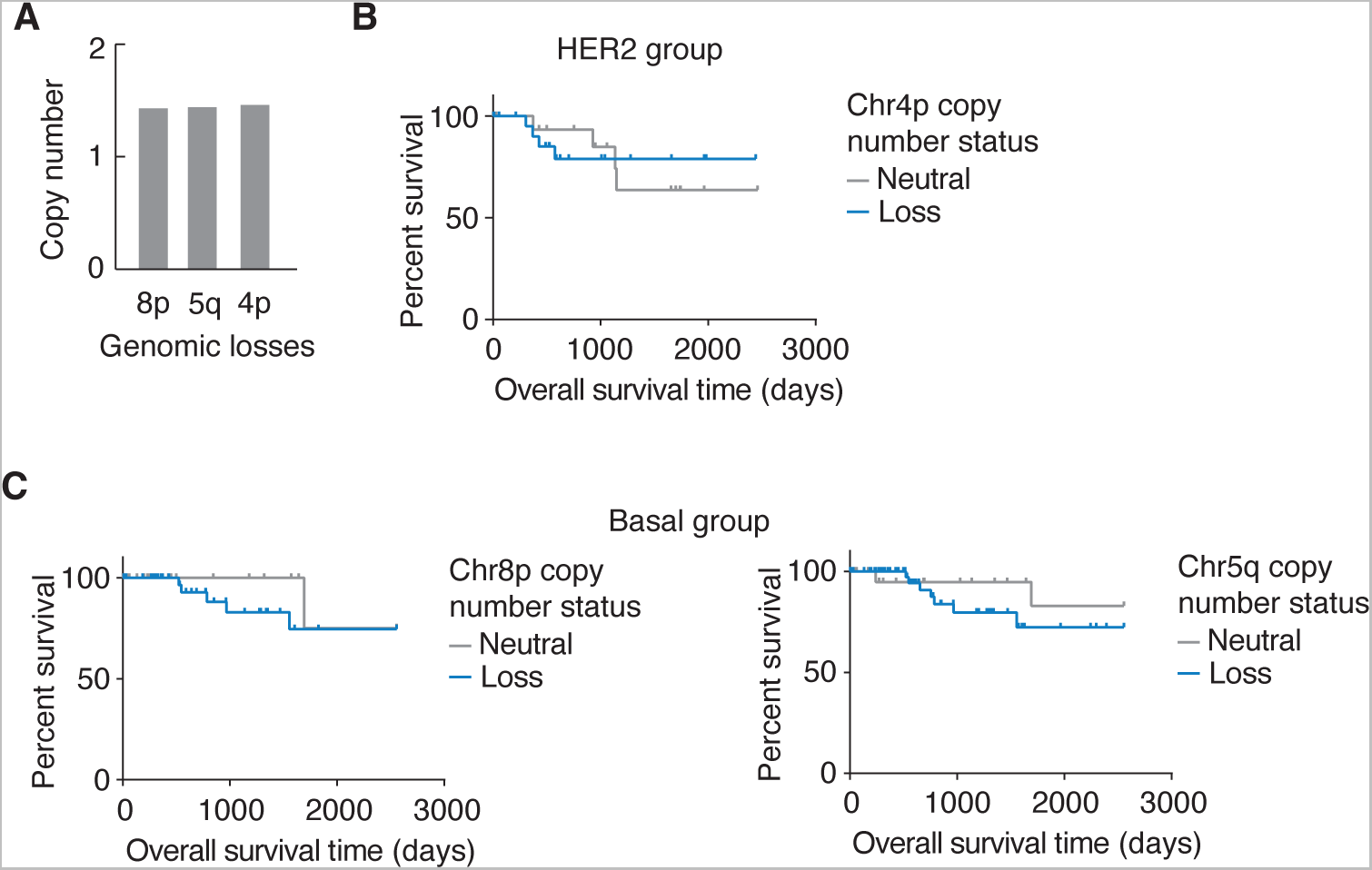
Recurrent large chromosomal deletions in breast cancer. **(A)** _Copy number was obtained from the TCGA segmented mean showing that frequently recurrent large chromosomal deletions in basal breast cancer are hemizygous, n = 91. **(B)** Overall survival of Her2 breast cancer patients with copy neutral and deletion status of chr4p shows no difference between groups, n = 55. **(C)** Overall survival of basal breast cancer patients with copy neutral and deletion status of chr8p and chr5q shows a trends towards a worse survival of patients with these chromosome arm losses (chr 8p loss *p* = 0.5, chr5q loss *p* = 0.4 as assessed by long rank test; n = 91)._

**Figure S2.**
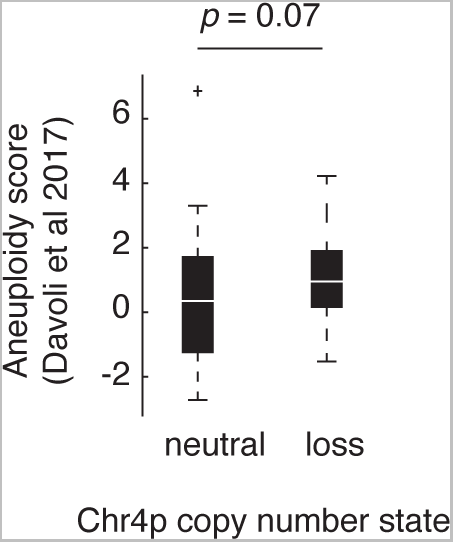
Aneuploidy in basal breast cancer with different chr4p copy number states. Aneuploidy score as quantified by Chrom.Arm.SCNA.Level median reported by Davoli et al Science 2017 shows no statistically significant difference in aneuploidy between chr4p copy neutral vs deletion basal breast cancer samples. Significance was assessed using Wilcoxon rank sum test.

**Figure S3.**
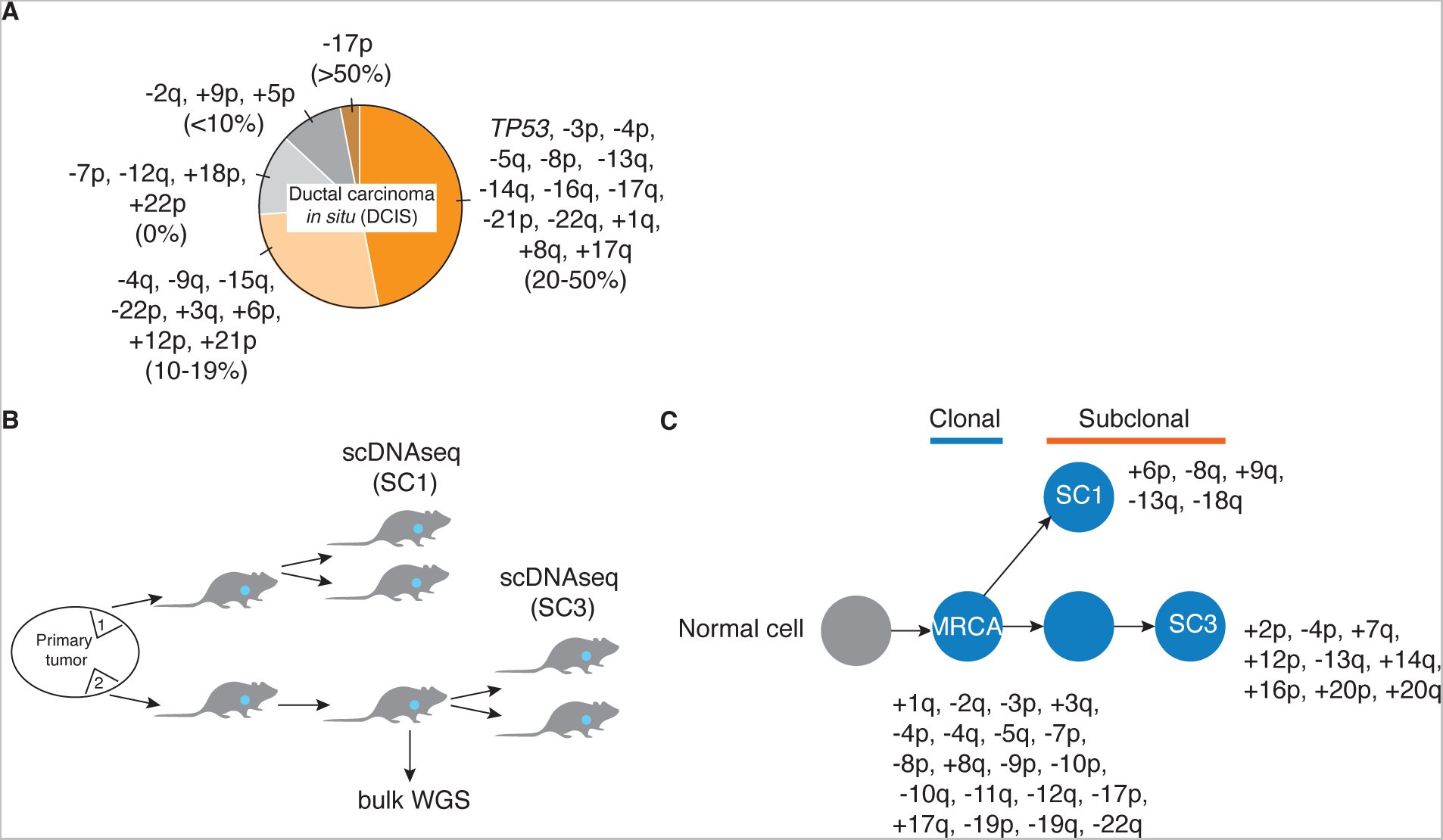
Frequency of genomic events in other datasets. **(A)** Genetic events from clonal and variable/constant regions of the timing analysis presented in Figure 2B were detected in Ductal Carcinoma *in situ* (DCIS) from a previous study (Lips et al *Nat Biotech* 2022). Frequency of patient samples from a total of 95 is shown in brackets. **(B)** Schematic of PDX generation, which was used for bulk WGS and scDNAseq. Single cell DNA sequencing was conducted on four GCRC1735 PDX samples. Two different locations within the primary tumor were biopsied, cryopreserved and propageted in NOD-SCID mice. Two mice were engrafted using a fragment derived from one location from passage 2 in the PDX and two mice were engrafted using a fragment drived from another location from passage 3 in the PDX. **(C)** CHISEL (Zaccaria et al *Nat Biotech* 2020) was used to generate an evolutionary timeline showing that chr4p loss is an early event in basal breast cancer PDX progression.

**Figure S4.**
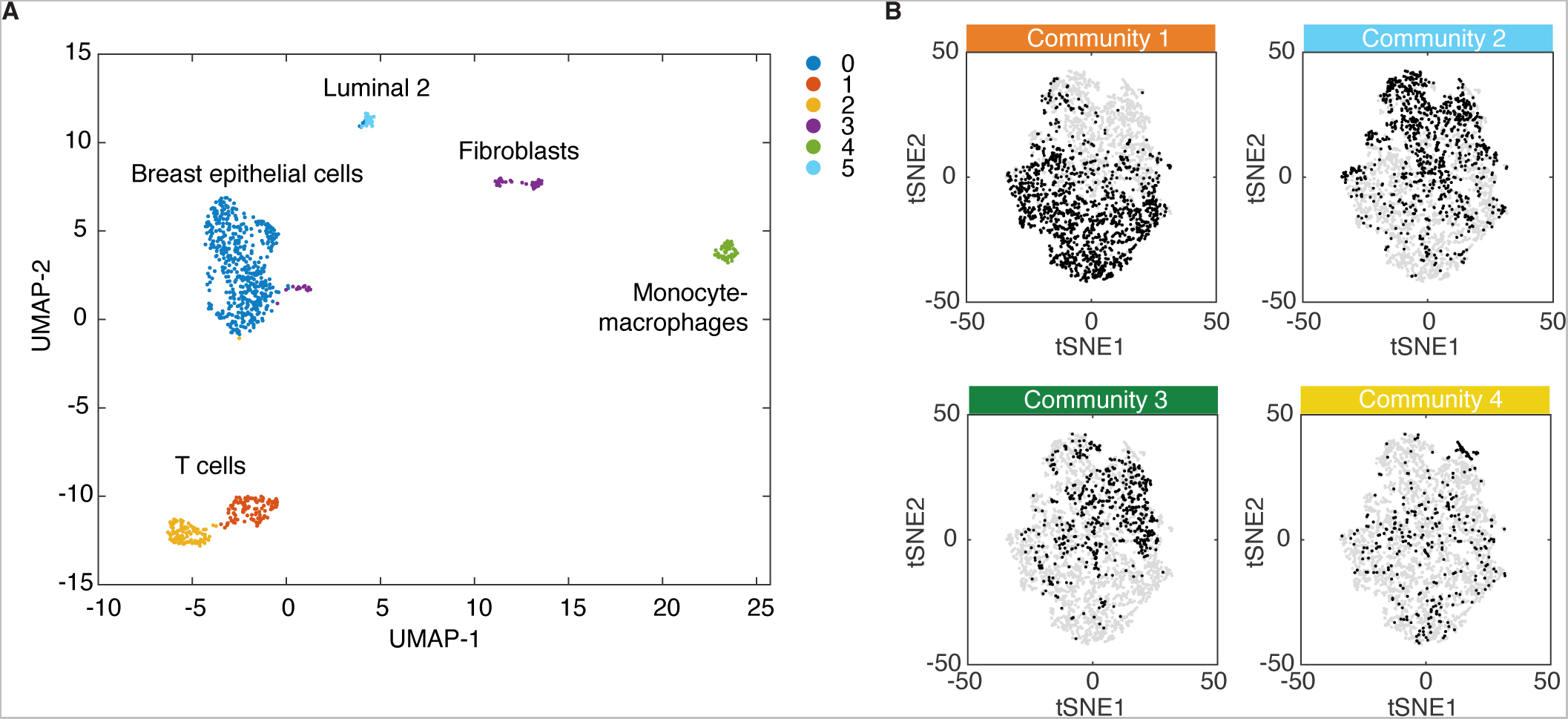
Inferring copy number aberrations from scRNAseq. **(A)** scRNAseq of normal breast epithelial cells. UMAP plot of single cell RNA seq data from two reduction mammoplasty samples. Clusters were annotated using previously defined cell type markers. **(B)** Overlapping inferred copy number communities with scRNAseq gene expression clusters. scRNAseq map as reported in a previous study is used to annotate single cells with inferred copy number communities as analyzed in this study. Single cells coloured in black belong to the specifiied community.

**Figure S5.**
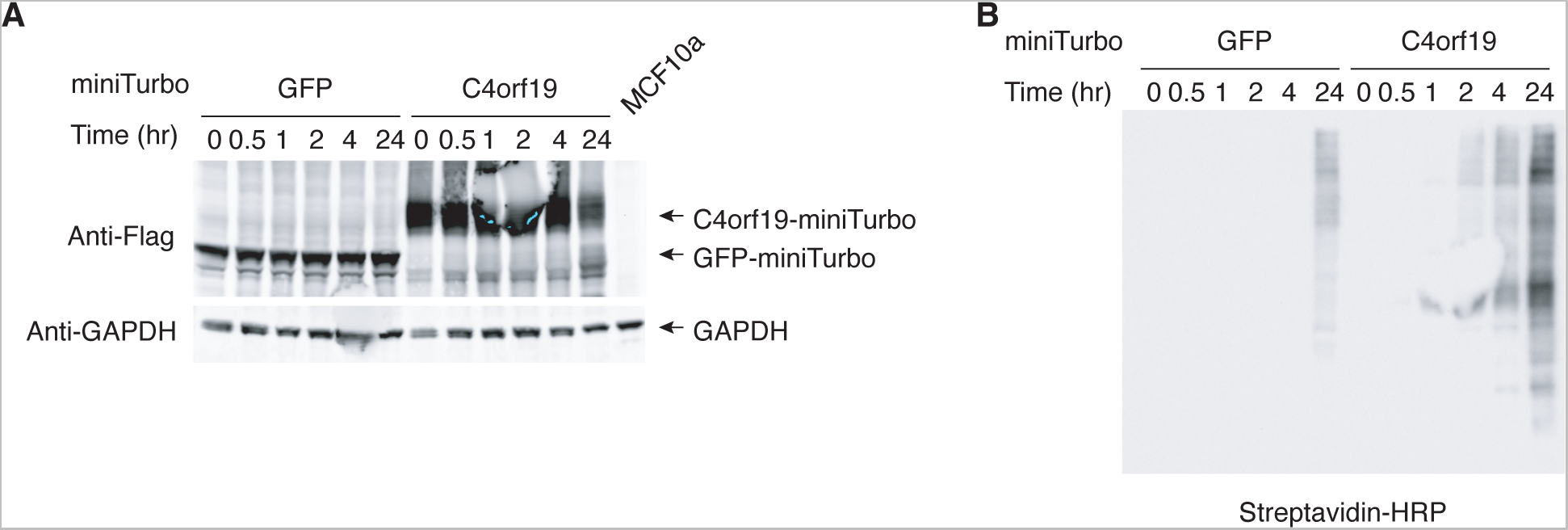
miniTurboID screen bait protein expression and biotinylation level. **(A)** miniTurboID C4orf19 protein bait expression in MCF10a. MCF10a cells stably expressing miniTurboID bait proteins: C4orf19-3XFLAG-miniTurbo and GFP-3XFLAG-miniTurbo control were induced with 0.5 µg/ml doxycycline for designated time. Protein bait expression was assessed using Anti-Flag antibody. GAPDH protein level served as the loading control. **(B)** Biotinylation level. Biotin labeling was induced with 0.5 µg/ml doxycycline and 40 µM biotin for designated time. Optimal biotinylation was achieved 4 hr post induction.

## Notes

### Competing Interest Statement

The authors have declared no competing interest.

### Summary of Updates

Revised Figure S3.

